# Hidden-state inference aligns attention and neural representational geometry during flexible behaviour

**DOI:** 10.1101/2025.10.10.681143

**Authors:** Casper Kerrén, Stephanie Theves, Mikael Johansson, Peter Gärdenfors, Christian Doeller

**Affiliations:** Max Planck Institute for Human Cognitive and Brain Sciences, Leipzig, Germany; Max Planck Institute for Empirical Aesthetics, Frankfurt, Germany; Department of Psychology, Lund University, Lund 221 00, Sweden; Cognitive Science, Lund University, Lund 221 00, Sweden; Kavli Ins,tute for Systems Neuroscience, Centre for Neural Computation, Egil and Pauline Braathen and Fred Kavli Centre for Cortical Microcircuits, Jebsen Centre for Alzheimer’s Disease, NTNU Norwegian University of Science and Technology, Trondheim, Norway

## Abstract

Flexible decision-making in uncertain environments requires inferring latent task states and prioritising behaviourally relevant information. We tested the hypothesis that this process is associated with systematic changes in attention and neural representational structure. Human participants performed a serial reversal learning task that required identifying which stimulus dimension(s) were currently relevant. Behaviourally, participants rapidly adapted to context switches, and a hidden-state inference model best explained these adaptations, outperforming multiple reinforcement-learning variants in predicting both choices and inferred contexts. Oculomotor behaviour provided converging evidence for selective attention, with gaze increasingly concentrated on relevant features, tracking trial-by-trial belief updates, and transiently broadening following high prediction errors. Guided by these results, we used magnetoencephalography to examine how latent-state inference was reflected in neural activity. Neural representations encoded the currently inferred context and were modulated by recent prediction errors, with larger prediction errors associated with weaker context representations on subsequent trials. Representational analyses further revealed that successful performance was associated with a systematic reorganisation of neural geometry, characterised by selective amplification of task-relevant representational distances and increased separation of behaviourally relevant stimulus values. Together, our findings demonstrate that latent-state inference, attention, and neural representational geometry are tightly coordinated during flexible decision-making, providing a systems-level account of how behaviourally relevant information is selectively prioritised and represented to support adaptive behaviour.

## Introduction

Human beings inhabit environments rich in diverse streams of information, yet the limited capacity of the cognitive system makes it impossible to process all inputs simultaneously. As a result, individuals must prioritise and selectively process information, flexibly adjusting behaviour in accordance with current goals. This adaptive capacity is especially apparent in decision-making, where humans and animals navigate complex environments with remarkable ease, often where classic reinforcement learning (RL) algorithms struggle (Bellman, 1957).

Rather than learning about all possible stimulus-outcome associations, it is often more efficient to represent the environment in a lower-dimensional manifold by focusing on task-relevant features (Konidaris, 2019; Niv, 2019). For example, a cost-conscious car buyer may focus on price and weight, whereas a performance-oriented buyer may prioritise speed, illustrating how contextual goals determine the processing of relevant features. This selective focus is thought to arise through representation learning, which mitigates the curse of dimensionality in RL by identifying a representational subspace aligned with current task goals (Bellman, 1957; Gershman & Niv, 2010; Leong et al., 2017; Niv, 2019; Sutton & Barto, 2018.; Wilson & Niv, 2012). Attentional filters are central to this process (Edgell & Morrissey, 1987; Kruschke, 1996; Nosofsky, 1986), enhancing dimensions deemed relevant by recent experience and internal goals. These filters, measurable via oculomotor behaviour (Blair et al., 2009; Corbetta & Shulman, 2002; Kowler et al., 1995; Moore & Fallah, 2001; Rehder & Hoffman, 2005), dynamically adapt as participants infer which features are currently relevant, providing a behavioural window into how inferred task structure guides attention. Through this mechanism, internal models are thought to interact with attention to support flexible, goal-directed behaviour (Bar-Gad et al., 2003; Corbetta & Shulman, 2002; Frank & Badre, 2012; Leong et al., 2017; Niv et al., 2015).

However, classic model-free RL approaches offer limited explanatory power in environments where reward contingencies shift or depend on unobservable variables. In contrast, inference-based approaches such as structure learning posit that agents form and update beliefs about latent states of the environment (Boyen et al., 2013; Braun et al., 2010; Gershman et al., 2015). Recent theoretical accounts further propose that such latent-state inference may be implemented via schema-like representations, which provide structured priors that guide reinforcement learning and generalisation across contexts (Bein & Niv, 2025). These beliefs are thought to integrate sensory observations with prior knowledge about the task’s causal structure, allowing for inference-guided behaviour under uncertainty. Thus, behaviour is shaped not only by reinforcement history, but also by inferences about hidden causes (Daw et al., 2011; Mishchanchuk et al., 2024; Niv et al., 2015; Starkweather et al., 2018; Vertechi et al., 2020; Wilson et al., 2014). A key distinction between latent-state inference paradigms and more traditional rule-switching tasks is that, in the latter, the relevant feature or rule is explicitly specified, whereas in the former it must be inferred under uncertainty from feedback. Even when behaviour converges to a stable strategy, latent inference requires maintaining and updating beliefs about task structure on a trial-by-trial basis. This distinction provides a unique opportunity to examine how belief dynamics are reflected in attention and neural representations, beyond what can be observed in explicitly instructed paradigms.

Neuroscientific evidence increasingly supports the idea that belief-guided dimensionality reduction underlies flexible behaviour. To manage environmental complexity, the brain appears to compresses high-dimensional input into lower-dimensional subspaces aligned with task goals (Behrens et al., 2018; Bellmund et al., 2018; Eichenbaum & Cohen, 2014; Garvert et al., 2017; Knudsen & Wallis, 2021; Leong et al., 2017; Mack et al., 2020; Menghi et al., 2021, 2023; Park et al., 2020; Radulescu et al., 2019; Schuck et al., 2016; Theves et al., 2020; Zhang et al., 2023; Zhang et al., 2024). Within these subspaces, attention is associated with systematic changes in representational geometry, enhancing relevant features while suppressing irrelevant ones. This process involves modulating representational axes, expanding dimensions aligned with current goals and compressing those that are not (Duan et al., 2024; Flesch et al., 2022; Kruschke, 1996; Love et al., 2004; Nosofsky, 1986; Park et al., 2023). Although such transformations are typically observed after extensive training (e.g., Flesch et al., 2022; Park et al., 2023) and are modulated by what is shown on screen (Duan et al., 2024), recent work in non-human primates suggests they can also emerge dynamically (Zhang et al., 2023), supporting flexible behaviour in volatile environments. Here, we extend this line of work by testing whether similar transformations are associated with latent-state inference, allowing us to examine how belief updating relates to changes in representational geometry in real time.

In the present <u>pre-registered</u> study, we asked whether participants could dynamically retrieve from memory the relevant latent subspace (context) and flexibly engage task-relevant aspects of the representational space on a trial-by-trial basis. To this end, we designed a serial reversal learning task in which participants judged cars based on three feature dimensions, each with five discrete levels (Fig. 1), controlling for psychological distance in a separate task (Supplementary Fig. S1). Crucially, while the stimulus space remained fixed, the task-relevant dimension varied according to an unobservable contextual variable. Participants were instructed to select cars for three different characters, where the value of a car depended on character-specific rules defined over one or two feature dimensions. Feedback was provided on correct trials, allowing participants to infer the currently relevant dimension through experience. By holding the stimulus space constant while varying task goals, the task enabled us to examine goal-directed representation learning.

**Figure 1.**
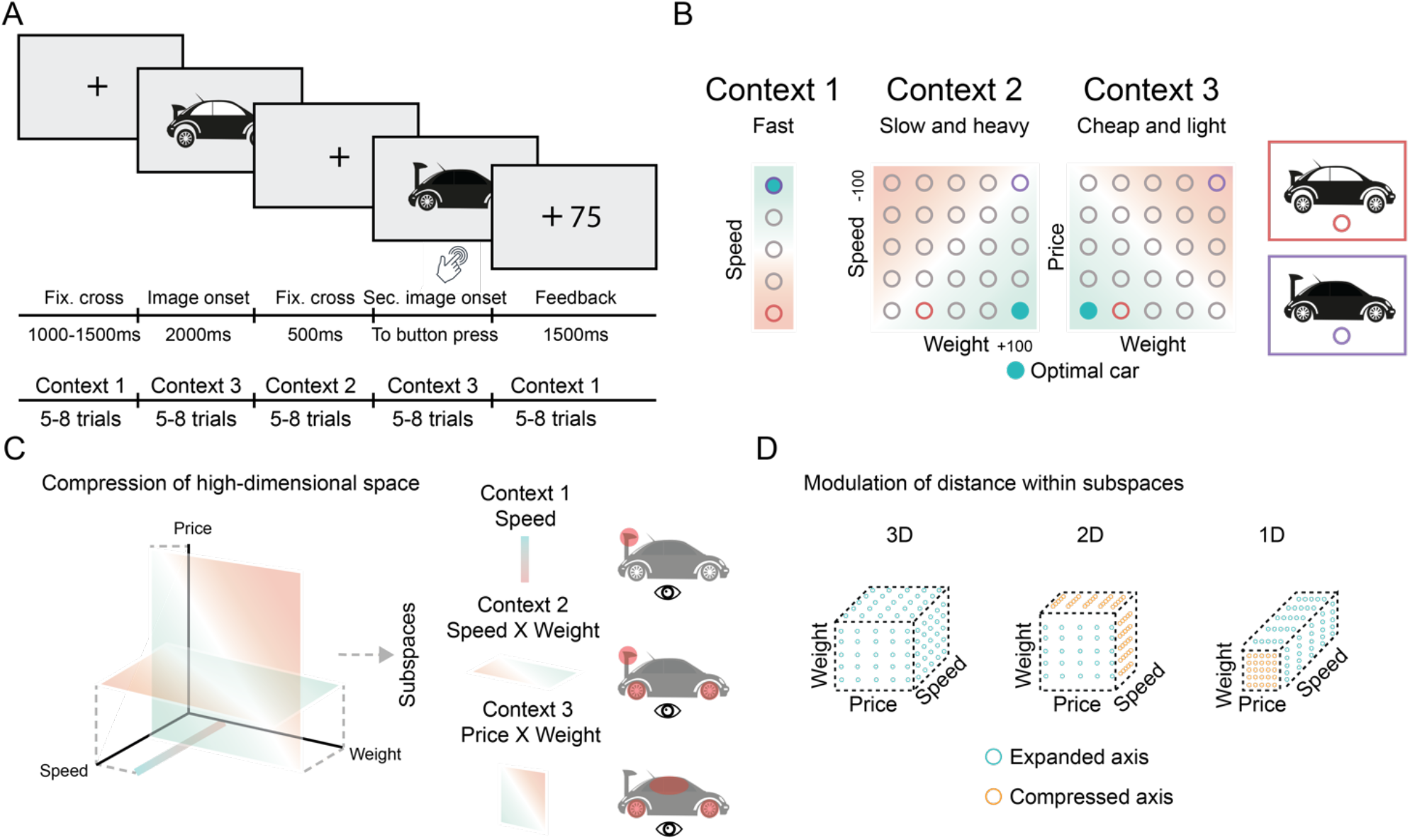
Context-dependent decision-making task. **A**. Each trial involved the presentation a fixation cross, one car, followed by another fixation cross and a second car. Participants were instructed to indicate which of the two cars had the higher value given the character (context) they thought they were playing. Timeline of a trial and rationale for context switches shown underneath. **B**. The reward values were defined according to three different stimulus-reward contingency maps. An example of three maps is shown, which across participants were counterbalanced. **C**. Hypothesised compression of a 3D space into subspaces for the different characters participants played. Green to orange gradient showing actual reward in a given context. Cars highlight the relevant dimensions, assessed with eye-tracking. **D**. Neurophysiological hypotheses propose that in an initially full representational space (3D), the distance between levels would be equal across axes, whereas when participants focused on relevant subspaces (2D or 1D), the axes within these subspaces would be expanded (relative to irrelevant ones; shown as green and orange circles, respectively).

To characterise the processes underlying flexible, context-sensitive behaviour, we first analysed participants’ behavioural performance to assess their ability to adapt to latent context switches. We then used computational modelling to adjudicate between candidate learning mechanisms, comparing reinforcement learning models to a hidden state inference model that tracks beliefs over unobservable task states. To evaluate whether such belief dynamics guided perceptual processing, we analysed oculomotor behaviour to test whether attention selectively converged on task-relevant features and further related eye-correlates to the winning model. Finally, we used magnetoencephalography (MEG) to examine whether neural representational dimensionality and geometry covaried with belief dynamics, specifically whether activity was differentially expressed when participants made the correct choices as compared to incorrect.

## Results

### Participants rapidly adapted behaviour following latent context switches

We first characterised overall task performance by examining reaction times (M = 1.14 secs, SE = .06; Fig. 2A) and accuracy (M = .77 secs, SE = .06; Fig. 2B) across trials. We next examined how performance evolved within each context block by computing reaction time and accuracy as a function of trial position (1-8; Fig. 2C-D). We expected reaction times to decrease and accuracy to increase across trials as participants adapted to the current context. To test these predictions, we fitted a generalised linear model (GLM) to each participant’s reaction times and accuracies, and evaluated the resulting slopes using Wilcoxon signed-rank tests (signrank function in MATLAB) against a null hypothesis of zero slope. Consistent with our hypotheses, we observed a significant negative slope for reaction time (z = -5.90, p < .01; Fig. 2C) and a positive slope for accuracy (z = 6.05, p < .01; Fig. 2D).

**Figure 2.**
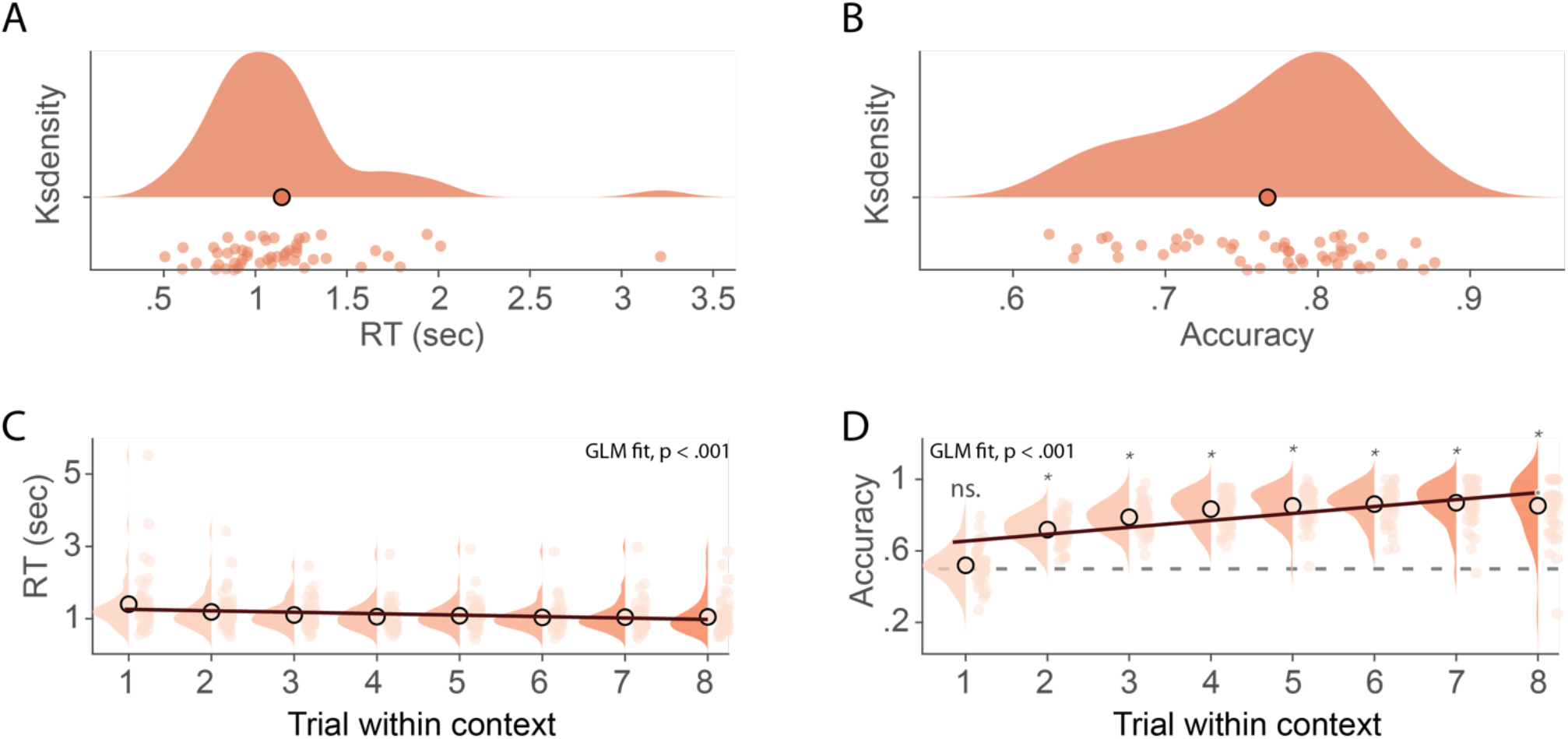
Behavioural results. **A**. Average reaction times (RTs) across all trials in the serial reversal learning task, with a mean of approximately 1.14 seconds. **B**. Average accuracy across all trials in the serial reversal learning task, with a mean of approximately 77%. **C**. Reaction time plotted as a function of trial position within a context block. A significant negative slope was observed, showing that reaction time decreased across trials (z = -5.90, p < .01). **D**. Accuracy plotted as a function of trial position within context blocks. A significant positive slope was observed, indicating improved performance with increasing evidence about the current context (z = 6.05, p < .01). Only the first trial in a new block showed no significant difference from chance (ns.).

To assess whether participants were sensitive to latent contextual structure, we compared accuracy on each trial to chance level (50%), based on the fact that participants were informed at the end of each context block that a switch was about to occur. As expected, performance was at chance only on the first trial following a switch (t(49) = 1.32, p = .19; uncorrected), suggesting that participants used feedback to rapidly adjust behaviour following context switches. Together, these results indicate that participants rapidly adapted behaviour following context changes and were sensitive to underlying task structure.

### Participants’ behaviour is best captured by hidden-state inference model

To characterise the computational processes underlying this adaptive behaviour, we compared candidate learning models that varied in how they handled latent structure. Specifically, we asked whether participants’ performance was best explained by reinforcement learning (RL) strategies, which update stimulus-reward values based on experienced outcomes, or by a hidden state inference (HSI) strategy, which maintains probabilistic beliefs over unobservable contexts.

On each trial, participants were required to determine which of three contexts was currently active, using only outcome feedback and cues indicating when a switch was about to occur. On the first trial after a switch, chance performance was expected (Fig. 2D). To learn over time, participants’ behaviour could, in principle, be accounted for by two broad classes of strategies: an RL approach, such as Q-learning, which updates value estimates incrementally based on observed outcomes, or a HSI approach, which constructs and maintains beliefs over a latent context variable that guides both choice and attention.

The four computational models (see Table S1 for model parameters) differed in how they update stimulus-reward associations and whether they represent hidden states explicitly. First, standard Q-learning (Q) provides a baseline incremental reinforcement learning strategy, in which action values are updated only based on experienced outcomes (Sutton & Barto, 2018; Watkins & Dayan, 1992). Second, Q-learning with forgetting (QF) extends this model by allowing the values of unchosen options to decay over time, capturing a potential recency bias or memory limitation (Kato & Morita, 2016). Third, Q-learning with counterfactual updating (QC) incorporates learning from unchosen outcomes by updating the values of both chosen and unchosen options based on their expected outcomes, thus allowing the model to simulate learning via exploration or internal evaluation (Palminteri et al., 2015). Finally, the HSI model assumes that participants infer which latent context is currently active, maintaining and updating probabilistic beliefs over these contexts based on feedback and cues. This model has been shown to account for rodent behaviour in similar tasks with latent reversals (Mishchanchuk et al., 2024) and formalises the idea that participants may rely on inferred task structure rather than relying solely on incremental value updating.

To model choices, we used context-specific stimulus-reward contingency matrices and marginalised over the three possible contexts, weighted by their posterior probabilities. Choice stochasticity was captured via a softmax with a free inverse temperature parameter (β_action). If a participant’s belief was weighted toward the correct context, the model was more likely to predict the correct action.

Model fits were evaluated using the Bayesian Information Criterion (BIC). The HSI model provided the best fit to the data (M = 308.10, SE = 6.38), followed by the counterfactual Q-learning model (QC; M = 464.78, SE = 7.73), the Q-learning with forgetting model (QF; M = 526.08, SE = 10.53), and standard Q-learning (Q; M = 495.64, SE = 11.80). The HSI model outperformed QC by a wide margin (t(49) = -21.70, p < .01) and was the best-fitting model for all participants (Fig. 3A).

**Figure 3.**
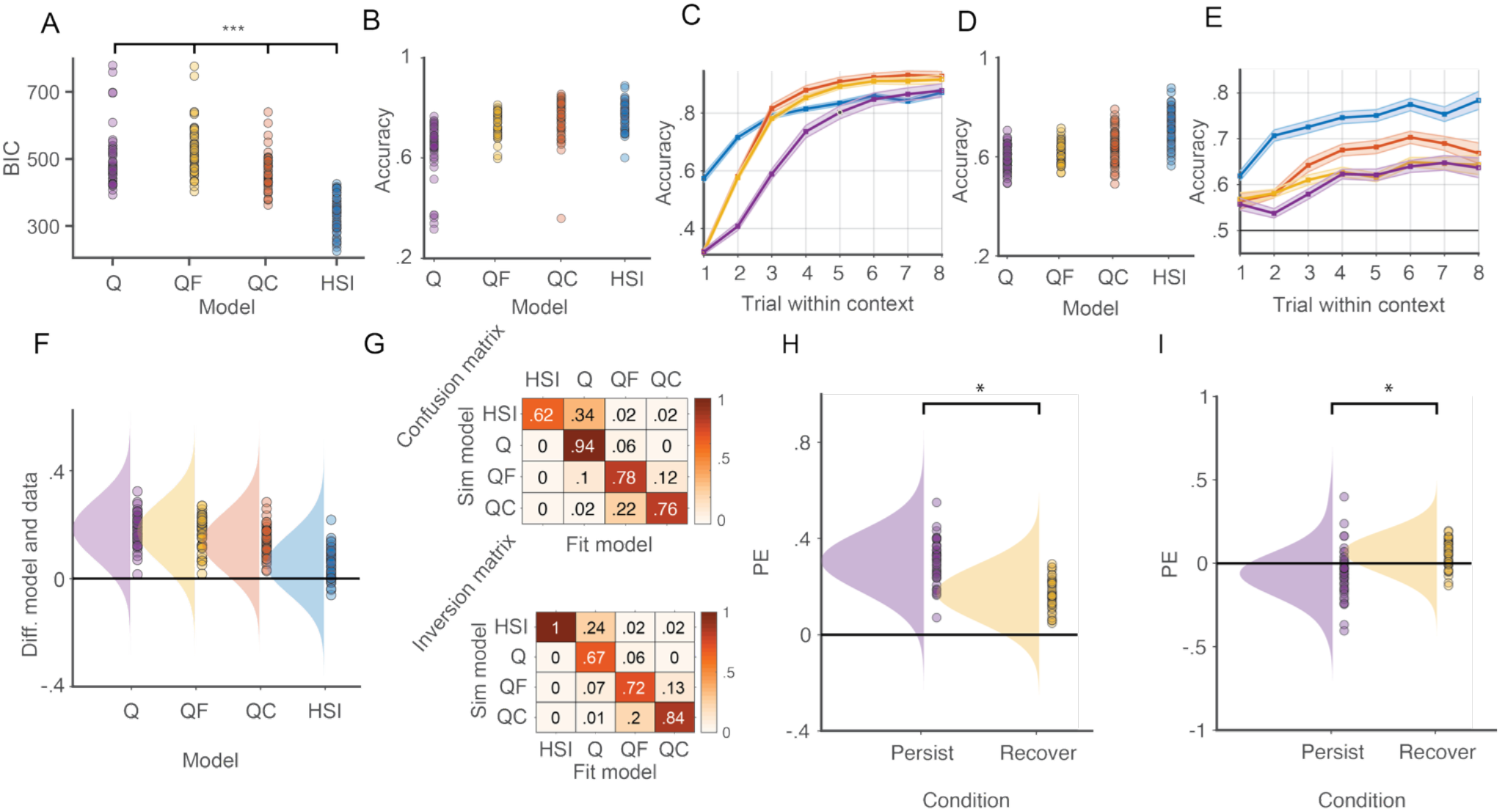
Hidden state inference model best fit the behavioural data. **A**. Bayesian information criterion (BIC) for Q-learning (Q), Q-learning with forgetting (QF), Q-learning with counterfactual updating (QC) and Hidden state inference model (HSI) showed significantly lower BIC scores for HSI compared to the other models, indicating that the HSI model showed the best fit to the data. **B**. Average of fraction of correctly predicted contexts for the four models. **C**. Same as (B), but now showing for each trial within a context. **D**. Average of fraction of correctly predicted button presses for the four models. **E**. Same as (D), but now showing for each trial within a context. **F**. Comparing predicted button presses to actual button presses showed that HSI was most aligned with actual data. **G**. We assessed model recoverability and identifiability by simulating data from each model and then fit it with each model. Highest values along the diagonal indicates that each model could best recover its own simulated data (top) and that the best-fitting model typically matched the generative model (bottom). **H**. Evaluating the model-based prediction error (PE) on trials following an initial error. When the model persisted to be wrong on the following trial, unsigned PE was significantly higher compared to when it recovered and predicted the right action. **I**. When instead evaluating the signed PE, we found that when the model recovered on the following trial, the PE was mostly positive, significantly more so compared to when it continued to predict the wrong action, where we saw mainly negative PE.

The HSI model also achieved the highest accuracy in predicting context (M = 77.46%, SE = 8.0%), compared to the second-best model QC (M = 75.35%, SE = 11.5%), QF (M = 73.68%, SE = 6.8%), and Q (M = 63.72%, SE = 15.2%). However, the difference between HSI and QC in predicting context was not statistically significant (t(49) = 1.55, p = .13; Fig. 3B-C).

A similar pattern was observed for predicting participants’ actions. The HSI model again achieved the highest accuracy (M = 74.56%, SE = 8.3%), followed by QC (M = 64.35%, SE = 9.5%), QF (M = 61.13%, SE = 6%), and Q (M = 59.70%, SE = 7%). The HSI model predicted actions significantly better than the QC model (t(49) = 18.06, p < .01), with all participants showing higher action prediction accuracy under HSI (Fig. 3D-E).

We also compared model predictions of action to actual behavioural data. The smallest discrepancy between predicted and observed actions was found for the HSI model (M= 3.23%, SE = 1.39%), followed by QC (M = 13.67%, SE = 1.58%), QF (M = 16.95%, SE = 1.63%), and Q (M = 18.19%, SE = 1.66%; Fig. 3F).

To confirm model identifiability, we conducted a model recovery analysis (Wilson & Collins, 2019) by simulating data with best-fitting parameters for each model and re-fitting all models to each dataset. The resulting confusion matrices showed strong diagonal structure (Fig. 3G), confirming that each model best recovered its own data and that the HSI model was reliably distinguishable from the RL-based models.

To further examine the relationship between model-derived belief dynamics and prediction error (PE), we computed trial-wise PE as the difference between observed choice accuracy and the model’s predicted probability of a correct response. We analysed PE in two complementary ways. First, using unsigned PE to quantify surprise irrespective of outcome valence, we examined whether surprise decreased when the model successfully updated its beliefs following an initial misprediction (inference trials; Courellis et al., 2024). On trials where the model was incorrect on trial 1 (i.e., immediately after a context switch), PE on trial 2 was significantly lower when the model recovered and predicted correctly (M = .17, SE = .01) than when it remained incorrect (M = .30, SE = .01; t(49) = 9.75, p < .01; Fig. 3H). This indicates that PE tracked trial-by-trial changes in model-derived belief updating.

We repeated the analysis using signed PE to examine outcome valence. When the model recovered on trial 2, signed PE was significantly positive (M = .07, SE = .01; t(49) = 7.01, p < .01), indicating a better-than-expected outcome. In contrast, when the model remained incorrect, signed PE was significantly negative (M = -.07, SE = .02; t(49) = -3.31, p < .01), reflecting a worse-than-expected outcome (Fig. 3I). These findings suggest that trial-wise PE reflected whether the model successfully updated its beliefs and encoded the valence of outcome evaluation, consistent with the idea that prediction error signals are related to belief updating and outcome evaluation during task performance.

Taken together, these results indicate that participants’ behaviour was best explained by a model-based inference framework, with the HSI model outperforming all alternatives in capturing both trial-by-trial choices and inferred contextual structure.

### Gaze selectively tracks model-derived task relevance

Next, we asked whether participants’ eye behaviour covaried with model-derived belief dynamics. If participants’ behaviour was guided by inferred context, this would be expected to bias attention toward the stimulus dimensions relevant for that context, and this, in turn, should be reflected in their eye movements. To assess this process, we analysed eye movements within predefined regions of interest (ROIs; the rear wing, window, and wheels; see Supplementary Fig. 2), which mapped onto either relevant or irrelevant features depending on whether the trial belonged to a 1D or 2D context block.

Previous studies have shown that gaze reliably tracks task-relevant dimensions during learning (e.g., Braunlich & Love, 2022; Leong et al., 2017; Rehder & Hoffman, 2005). We replicated and extended this finding: using gaze samples across trials within a context, the total number of attended dimensions decreased, converging toward the actual number of relevant dimensions by the final trials (Fig. 4A, left). We quantified this by segmenting each trial into an early window (from first image onset to fixation cross) and a late window (from second image onset to response). Participants attended to significantly fewer dimensions during the late window (M = 1.36, SE = .04) compared to the early window (M = 1.84, SE = .05; t(49) = 12.57, p < .01; Fig. 4A, right), consistent with a narrowing of attention toward task-relevant features.

**Figure 4.**
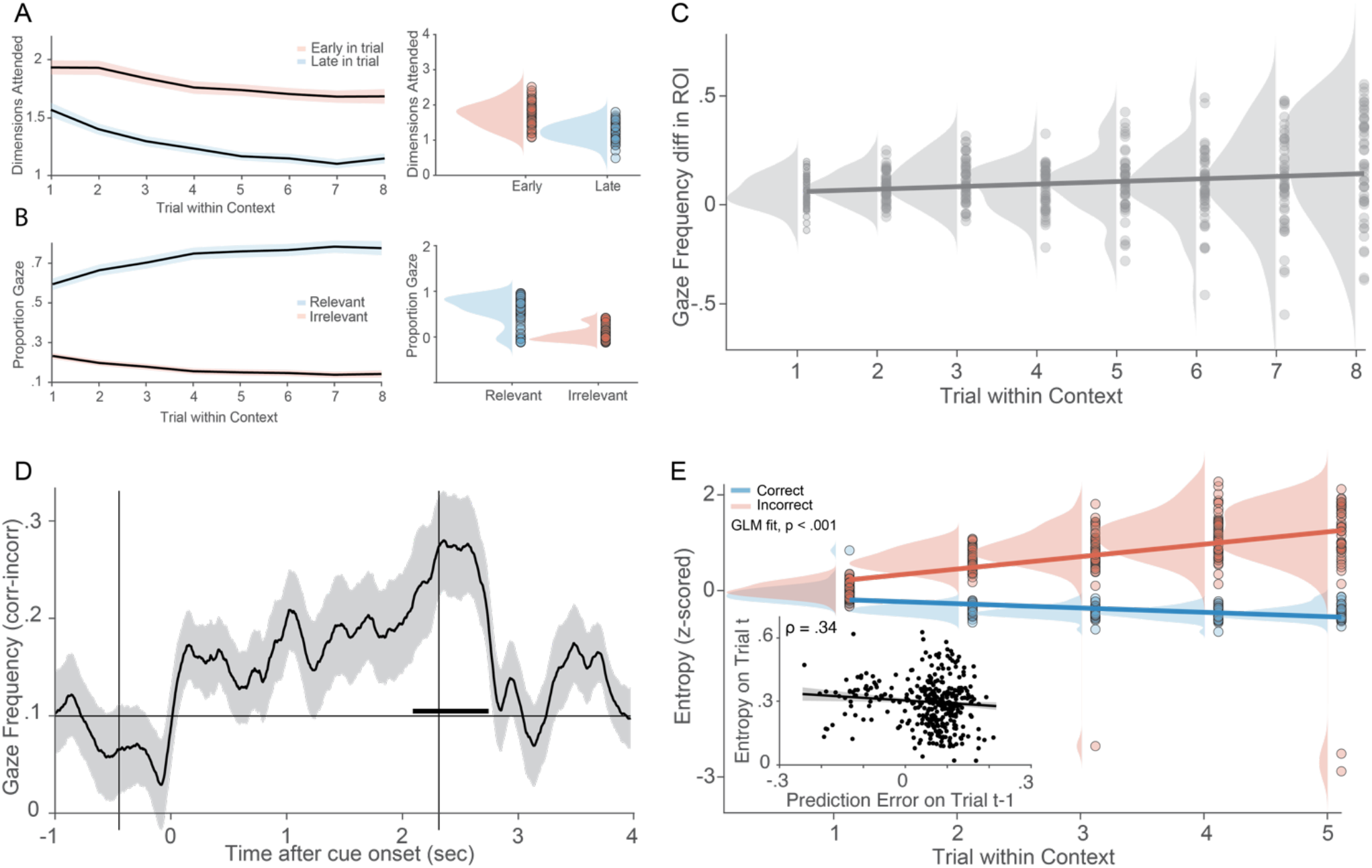
Context-guided eye-behaviour. **A**. Dimensions attended to for early (after first image onset to fixation cross; red) and late time points in trial (after second image onset to reaction time; blue), per trial within a context (left) and average across trials within contexts (right). At late time points participants focused more on the relevant dimensions (1 dimension on y-axis reflects 1 dimension for 1D trials and 2 dimensions for 2D trials). **B**. The proportion of gaze frequency for all trials between relevant (blue) and irrelevant (red) dimensions, shown for each trial within a context (left) and averaged across trials (right). Participants focused significantly more at the relevant dimensions as compared to the irrelevant dimensions **C**. Gaze frequency difference between correct and incorrect trials showed a significant positive slope, indicating that when participants where correct, their gaze was focused more within the relevant regions. **D**. When participants made an inference they shifted their gaze to the relevant ROI, as compared to when they did not make an inference, with a significance onset around second image onset. **E**. Entropy as a function of trial within a context. When participants were correct, entropy - a measure of disorganisation in gaze patterns reflecting how consistently across time within a trial participants focused on relevant features - significantly decreased across trials (blue), whereas when they were incorrect, entropy significantly increased (red). **Inset in E**. Signed PE predicted gaze entropy such that more negative PEs were related with more broadened visual sampling, whereas positive PEs were related to more focused attention.

To characterise how this attentional tuning evolved across trials, we computed the proportion of gaze samples landing on relevant versus irrelevant ROIs separately for each trial position (1-8) and participant, within the late time window. Overlapping features were counted as irrelevant only when they did not belong to the currently relevant feature set. These proportions were then averaged across participants and collapsed across 1D and 2D contexts. As shown in Fig. 4B (left), gaze allocation toward relevant features increased systematically with trial progression, while gaze to irrelevant features declined. Collapsing across trials, participants devoted significantly more gaze to relevant (M = .68, SE = .02) than to irrelevant features (M = .16, SE = .01; t(49) = 16.50, p < .01; Fig. 4B, right).

Next, we asked whether gaze to relevant features predicted correct performance. We fit a GLM to the difference in gaze allocation between correct and incorrect trials using gaze density measures from the late window. Wilcoxon signed-rank tests on model coefficients revealed significantly greater fixation within relevant ROIs on correct trials (z = 2.86, p = .004; Fig. 4C), indicating a relationship between attentional allocation and task success.

To probe this link further, we focused on inference trials. These trials are informative because they are associated with changes in model-derived beliefs about the relevant feature. We compared these trials to trials where participants were incorrect on both the first and current trial within a new context. A time-resolved analysis, corrected using cluster-based permutation testing, revealed a gradual increase in gaze toward the relevant dimensions starting at first image onset, with significant differentiation emerging around the second image onset (p_cluster_ < .05; Fig. 4D).

Finally, we assessed attentional uncertainty using gaze entropy, a measure of disorganisation in gaze patterns reflecting how consistently across time within a trial participants focused on relevant features (Shiferaw et al., 2019), with low entropy values reflecting more stable focus. We predicted that entropy would be highest immediately after a context switch and decline as participants inferred the relevant feature. We computed binary time series indicating whether gaze landed on relevant features and calculated entropy separately for correct and incorrect trials. After z-scoring entropy vectors, we fit GLMs to assess changes across trial position and tested slope values against zero. As predicted, entropy declined significantly across correct trials (z = -6.15, p < .01), while entropy increased across incorrect trials (z = 5.19, p < .01; Fig. 4E), suggesting that attentional stabilisation was associated with successful task performance and model-consistent belief updating.

Taken together, these results show that gaze allocation dynamically tracked task-relevant features in a manner consistent with latent structure, converging on the relevant dimensions as model-derived beliefs about context stabilised across trials. These findings demonstrate a close relationship between belief dynamics and attentional allocation, suggesting that inferred task structure is reflected in how visual information is selectively sampled during decision-making.

### Linking model-derived belief dynamics to visual attention

As a next step, we asked whether trial-by-trial variation in gaze behaviour covaried with model-derived belief dynamics. Specifically, if gaze is related to belief-guided attention, then trials with more focused gaze should be associated with stronger model predictions (i.e., higher certainty about the correct action according to the HSI model).

Gaze entropy was significantly lower on trials where the model correctly predicted the participant’s button press (M = .30, SE = .01) compared to trials where it did not (M = .33, SE = .01; t(49) = 4.05, p < .01), indicating that more focused visual attention was associated with behaviour that was better predicted by the model.

We next tested whether signed PE was associated with subsequent gaze behaviour. A linear mixed-effects model revealed that PE on trial t-1 significantly predicted gaze entropy on trial t (β = -.17, p = .02), such that more negative PEs were associated with more diffuse gaze distributions, while more positive PEs were followed by more focused attention (Fig. 4E, inset). Rather than selectively attending to a single feature dimension, participants showed broader attentional sampling following negative outcomes, consistent with exploratory behaviour and with changes in model-derived belief uncertainty.

Motivated by this link between belief dynamics and gaze, we then incorporated gaze metrics directly into the HSI model, allowing gaze entropy and gaze distribution to modulate belief updating and/or action selection. However, none of these augmented models outperformed the original HSI model in terms of BIC or choice prediction accuracy.

Together, these results demonstrate a close relationship between model-derived belief dynamics and attentional allocation. Crucially, this relationship is expressed at the level of trial-by-trial belief fluctuations under latent structure, rather than reflecting responses to explicitly instructed task demands. Gaze patterns systematically tracked model predictions, becoming more focused as belief certainty increased and more diffuse following negative outcomes. However, incorporating gaze into the model did not improve predictive performance, suggesting that both gaze and choice reflect shared underlying task representations rather than independent sources of information.

### Decoding of context is modulated by prediction error

Having established that behaviour and gaze were associated with model-derived belief dynamics, we next asked whether latent contextual representations could be decoded from neural activity and whether these representations were similarly modulated by PE. To address this, we trained multiclass classifiers to decode the currently active context from time-resolved neural activity patterns across electrodes. At each time point, classifiers were trained to distinguish among the three latent contexts using trial-wise neural features, yielding a continuous measure of neural evidence for the inferred context.

Across all trials, we found significantly above chance decoding accuracy shortly after first stimulus onset and remained elevated throughout the trial period with a peak shortly after second stimulus onset (cluster-corrected p < .05; Fig. 5A). When contrasting correct and incorrect trials, we found significantly more evidence for the correct class on correct than incorrect trials (cluster-corrected p < .05; Fig. 5B), with an early effect around 250ms after first image onset and during the later stages of the trial following presentation of the second stimulus. This indicates that successful behaviour was associated with more robust neural representations of the inferred latent context.

**Figure 5.**
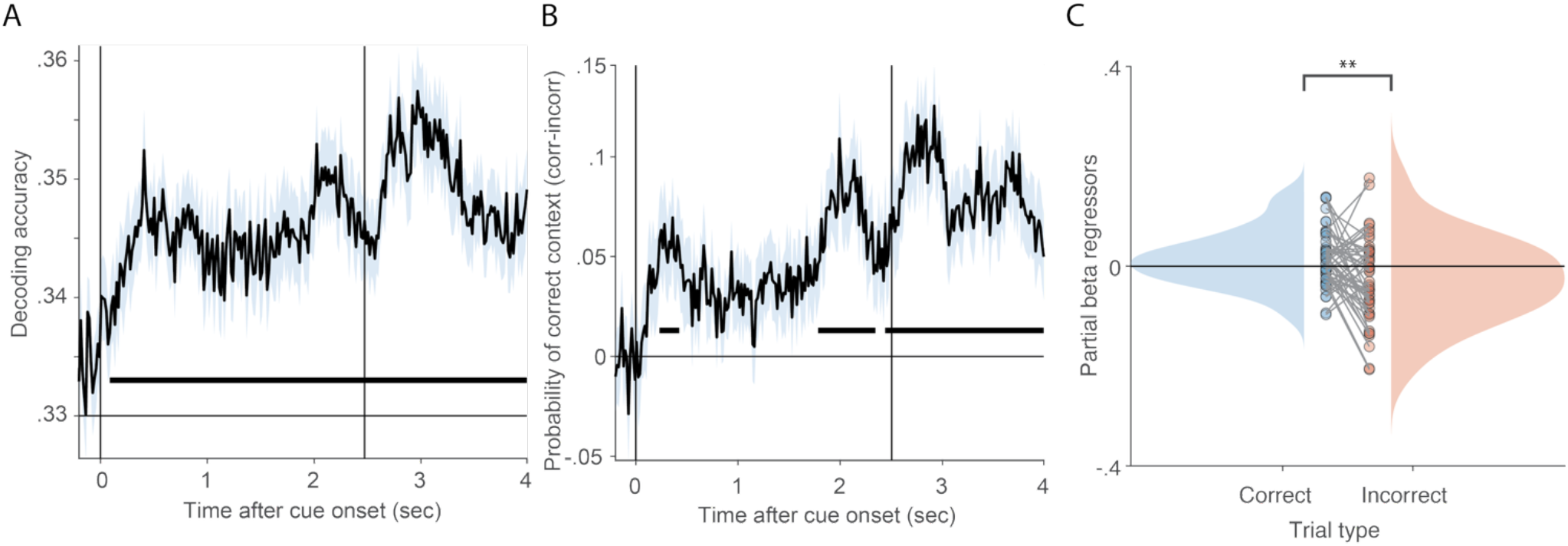
Decoding of context and frontal midline theta and PE. **A**. Time-resolved decoding of latent context from MEG activity revealed significant above-chance decoding beginning shortly after first stimulus onset and persisting throughout the trial. **B**. Neural evidence for the true context was significantly greater on correct than incorrect trials, with effects around 250ms after first image onset as well as shortly before second stimulus onset continuing until the end of the trial. **C**. Regression coefficients relating previous-trial signed prediction error (PE t-1) to current-trial neural context evidence differed significantly between correct and incorrect trials. Larger negative prediction errors on the preceding trial were associated with weaker subsequent context representations, whereas positive prediction errors were associated with stronger context representations, suggesting that negative outcomes destabilised and positive outcomes reinforced the inferred latent state.

To test whether trial-by-trial fluctuations in neural context evidence were related to model-derived belief dynamics, we fit a regression model predicting decoded neural evidence from belief strength, entropy, and signed prediction error. Guided by the results showing that prediction error on the previous trial was related to gaze entropy on the current trial, we used prediction error from the previous trial (PE t-1) while controlling for current-trial belief and entropy. Regression coefficients were estimated separately for correct and incorrect trials.

This analysis revealed a marked dissociation between correct and incorrect trials within the significant time window identified in Fig. 5B. On incorrect trials, larger negative PE t-1 values predicted weaker neural evidence for the current context (β = -.026, p = .028), whereas this relationship was weakly positive on correct trials (β = .015, p = .056). Direct comparison confirmed that the effect of PE differed significantly between correct and incorrect trials (t(49) = 3.09, p = .003; Fig. 5C).

Together, these findings suggest that neural representations of latent context are dynamically shaped by recent prediction errors. Negative prediction errors on the preceding trial were associated with weaker subsequent context representations, whereas positive prediction errors were associated with stronger context representations. This pattern is consistent with the idea that unexpected negative outcomes destabilise the currently inferred latent state, whereas unexpected positive outcomes reinforce it.

As a complementary validation of the HSI model-derived latent-state signals, frontal theta activity tracked both contextual uncertainty following context switches and trial-by-trial prediction errors generated by the model (Supplementary Fig. 3), consistent with previous work linking frontal theta to uncertainty monitoring and behavioural adaptation.

### Neural representational geometry differentially reflects successful and unsuccessful retrieval

Having characterised the overall dimensionality and its relationship to feature separability (Supplementary Fig. 4), we next asked whether the geometry of neural representations differed between correct and incorrect trials. Specifically, we tested whether successful trials were associated with a systematic reorganisation of representational distances along the task-relevant stimulus dimension as compared to unsuccesful trials.

We constructed a set of model representational dissimilarity matrices (RDMs; Fig. 6A; Supplemental Fig. 5-6) that captured distinct hypotheses about how stimulus representations might be organised. These included models reflecting a linear ordering of stimulus levels, an expansion of distances between extreme stimulus values, increased separation of extreme relative to intermediate values, salience-based coding, and centre-versus-edge coding.

**Figure 6.**
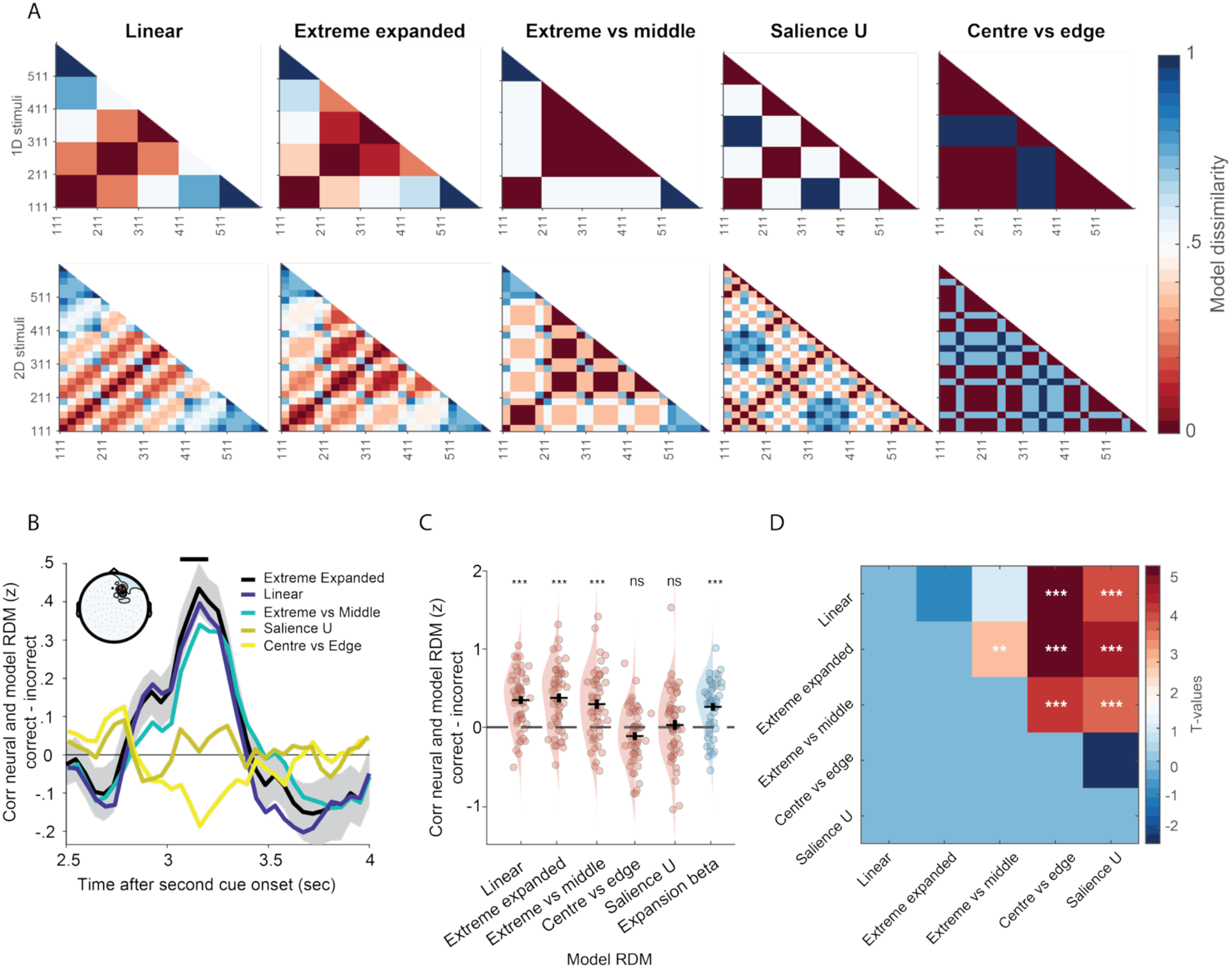
Successful retrieval is associated with an expanded task-relevant representational geometry. **A**. Model representational dissimilarity matrices (RDMs) used to characterise alternative representational geometries for 1D and 2D trials. For visualisation purposes, 1D models are shown for the case in which the first stimulus dimension is task-relevant, whereas 2D models are shown for the case in which the first and second dimensions are jointly relevant. Equivalent RDMs for all possible relevant dimensions are shown in Supplementary Figure 5-6. **B**. Correct-incorrect differences in neural representational similarity for all model geometries. Black line with grey shading shows cluster-based permutation analysis comparing neural representational similarity for correct and incorrect trials using the extreme-expanded model. A significant frontocentral cluster emerged approximately 500 ms after second-cue onset (black bar; topography inset). Shaded regions indicate ±SE across participants. Linear, extreme-expanded, and extreme-versus-middle models exhibited similar temporal profiles, whereas salience-U and centre-versus-edge models showed substantially weaker effects. **C**. Mean RSA values averaged across the significant cluster identified in panel B. Neural RDMs showed significant positive correspondence with the linear, extreme-expanded, and extreme-versus-middle models (FDR-corrected), but not with the salience-U or centre-versus-edge models. The expansion coefficient, obtained by regressing trial-wise neural distances against distances in the task-relevant stimulus space, was also significantly greater on correct than incorrect trials. Points represent individual participants; horizontal bars indicate group means ± SE. **D**. Pairwise comparisons of model fits within the significant cluster. Models capturing ordered stimulus structure (linear, extreme-expanded, and extreme-versus-middle) produced significantly larger correct-incorrect effects than salience-U and centre-versus-edge models, indicating that successful retrieval was associated with a representational geometry that preserved ordinal relationships while selectively increasing the separation of extreme stimulus values. Asterisks denote significance levels (*p < .05, **p < .01, ***p < .001).

Neural RDMs were computed by estimating the dissimilarity between activity patterns elicited during the first image (250-500 ms after onset) and activity patterns during the second image (2500-4000 ms after onset). The 250-500 ms window was selected based on the peak context decoding observed in Figure 5A and B, thus being unrelated to the current analysis. Neural and model RDMs were then compared using Spearman correlations. To determine whether observed correlations reflected meaningful representational structure rather than arbitrary similarity patterns, we generated participant-specific null distributions by randomly shuffling model geometries prior to correlation. This procedure was repeated 250 times for each participant.

Cluster-based permutation tests were performed separately for each model. To control for multiple comparisons across the five models, the cluster-forming significance threshold was adjusted to p = .01 (.05/5). Under this criterion, we observed a significant cluster for the extreme-expanded model (p = .006; Fig. 6B), indicating a stronger correspondence between neural and model geometries on correct relative to incorrect trials. No significant clusters were observed for the linear (p = .43), extreme-versus-middle (p = .20), salience-U (p = .03), or centre-versus-edge models (no significant clusters).

Within the significant cluster identified for the extreme-expanded model, correlations were significantly more positive than chance on correct trials (t(49) = 3.09, p = .003) and significantly more negative than chance on incorrect trials (t(49) = -5.58, p < .001). Thus, successful trials were associated with an ordered representational geometry in which task-relevant distances were amplified, with the strongest amplification observed for extreme stimulus values, whereas unsuccessful trials were associated with a reversal of this representational structure.

We next used the significant channels identified by the extreme-expanded cluster to examine the temporal evolution of the effect across the remaining models. Linear, extreme-expanded, and extreme-versus-middle models exhibited highly similar temporal profiles, whereas salience-U and centre-versus-edge models showed substantially weaker effects (Fig. 6B). Averaging across the significant channels and time window revealed significant correct-incorrect differences for the linear (t(49) = 6.77, p < .001), extreme-expanded (t(49) = 6.46, p < .001), and extreme-versus-middle models (t(49) = 4.71, p < .001), but not for the salience-U (t(49) = 0.46, p = .65) or centre-versus-edge models (t(49) = -2.19, p = .97; Fig. 6C; one-sided tests, false-discovery rate corrected).

To provide a model-independent measure of representational expansion, we additionally regressed trial-wise neural distances against distances in the task-relevant stimulus space. Consistent with the RSA results, the resulting expansion coefficient was significantly greater on correct than incorrect trials (t(49) = 5.32, p < .001; Fig. 6C, blue), indicating that successful trials were associated with stronger scaling of neural distances according to task-relevant stimulus differences.

Finally, pairwise comparisons between model fits revealed significantly larger correct-incorrect effects for models capturing ordered stimulus structure (linear, extreme-expanded, and extreme-versus-middle) than for salience-U and centre-versus-edge models (Fig. 6D). Furthermore, the significant extreme-expanded cluster exceeded a shuffled-RDM max-cluster null distribution (p = .004), demonstrating that the observed effect could not be explained by generic similarity structure alone.

Together, these results indicate that successful trials are associated with a structured reorganisation of representational geometry that preserves ordinal relationships between stimulus levels while selectively increasing the separation of extreme task-relevant values. In contrast, incorrect trials were characterised by a weakening or reversal of this geometry, suggesting that accurate performance depends on the emergence of an expanded task-relevant representational space.

## Discussion

In uncertain and dynamic environments, adaptive behaviour depends on the ability to track latent structure and prioritise task-relevant information. Here, we show that participants rapidly adapted to context changes in a serial reversal learning task, with coordinated changes in behaviour, visual attention, and neural representations. Across these domains, performance was best captured by a hidden-state inference (HSI) model, and both gaze behaviour and neural signals covaried with model-derived dynamics. Together, these findings provide converging evidence that belief-based representations of task structure are closely linked to how information is selectively processed and represented during flexible decision-making (Niv et al., 2015; Wilson et al., 2014).

Participants quickly adapted to the current context, as evidenced by systematic improvements in accuracy and reaction time across within-context trials. An HSI model captured these behavioural adaptations more accurately than multiple RL variants by explicitly maintaining probabilistic beliefs over latent task states (Boyen et al., 2013; Braun et al., 2010). The HSI model not only predicted choices and inferred contexts more accurately than its RL counterparts, but also showed high model recoverability and participant-level identifiability (Fig. 3). We note that our task structure may favour inference-based strategies once participants had learned the limited set of possible contexts, such that reinforcement learning and hidden-state inference may reflect different regimes of behaviour across learning rather than mutually exclusive processes. That human behaviour aligned most closely with this structure-sensitive model, even in a task with three contexts and frequent switches, is consistent with the view that participants relied on structure-sensitive representations to infer latent structure rather than relying solely on incremental reward histories (Daw et al., 2011; Vertechi et al., 2020). These results parallel recent findings in rodents (Costa et al., 2015; Mishchanchuk et al., 2024; Vertechi et al., 2020), extending the cross-species evidence for inference-based accounts of adaptive control.

Crucially, these belief-related dynamics were also evident in participants’ eye movements. Over trials, gaze selectively converged on task-relevant dimensions, and this attentional narrowing was more pronounced on correct trials. Gaze entropy declined over time within context blocks, indicating an increasingly structured sampling policy consistent with increasing sensitivity to task-relevant context (Najemnik & Geisler, 2005; Nelson & Cottrell, 2007; Rehder & Hoffman, 2005; Yang et al., 2016; Fig. 4A-B). Moreover, trial-level fluctuations in gaze patterns covaried with model-derived belief dynamics: entropy was lower when the model correctly predicted behaviour and increased following high prediction errors, consistent with exploratory sampling in the face of uncertainty (Fig. 4E, inset). Importantly, this relationship reflects inferred rather than explicitly instructed task relevance, distinguishing the present findings from prior work using externally cued task demands. These findings suggest that visual attention systematically prioritised task-relevant features in a manner consistent with inferred relevance (Corbetta & Shulman, 2002; Moore & Fallah, 2001).

While attention and model-derived beliefs were closely aligned, their relationship does not imply a specific mechanistic dependency. Incorporating gaze measures into the HSI model did not improve predictive performance, suggesting that gaze and choice were both systematically related to the same task structure captured by the model, rather than providing independent sources of information. In this sense, attention appears tightly coordinated with belief dynamics without specifying whether it is upstream, downstream, or part of the same underlying process (Bar-Gad et al., 2003).

As a complementary validation of the model-derived latent-state signals, frontal midline theta activity tracked both contextual uncertainty and model-derived prediction errors (Supplementary Fig. 3), consistent with previous work linking theta oscillations to belief updating, uncertainty monitoring, and outcome evaluation (Cavanagh & Frank, 2014; Cavanagh & Shackman, 2015; Cockburn & Holroyd, 2018; Heydari & Holroyd, 2016; Lopez-Gamundi et al., 2024). These findings indicate that neural signatures classically associated with cognitive control and reinforcement learning are also sensitive to prediction errors estimated from an inference-based model of latent task structure.

Beyond these outcome-related signals, we found direct evidence that latent-state representations were expressed in neural population activity. Context could be decoded from MEG activity shortly after stimulus onset and remained represented throughout the trial, with stronger context evidence observed on correct than incorrect trials. Importantly, recent prediction errors influenced these neural representations. Larger negative prediction errors on the previous trial predicted weaker neural evidence for the current context on subsequent incorrect trials, whereas positive prediction errors were associated with marginally stronger context evidence. Together with the gaze entropy findings, this suggests that prediction errors do not merely signal discrepancies between expected and observed outcomes, but may influence the stability of the inferred latent state itself. Negative prediction errors appear to destabilise latent-state representations, promoting broader attentional sampling and less reliable contextual representations on subsequent trials, whereas positive prediction errors may reinforce the currently inferred state.

Consistent with this interpretation, supplementary analyses showed that the effective dimensionality (ED) of neural activity scaled with task demands, with 1D models best accounting for 1D trials and both joint 2D and independent 1D representations contributing to performance on 2D trials (Supplementary Fig. 4A). Dimensionality was higher on correct than incorrect trials and positively tracked stimulus separability (Supplementary Fig. 4B-C), indicating that successful performance was associated with richer and more discriminable neural representations. These findings extend prior demonstrations of dimensionality compression during learning (Ahlheim & Love, 2018; Mack et al., 2020) by showing that representational dimensionality varied dynamically with task demands and behavioural performance.

Most importantly, we found that successful retrieval was associated with a systematic reorganisation of neural representational geometry. Neural population activity was best explained by geometries that preserved ordinal relationships between stimulus values while selectively amplifying distances along task-relevant dimensions. This effect emerged shortly after presentation of the second stimulus and was strongest for models in which extreme stimulus values were disproportionately separated (Fig. 6). Importantly, these effects were observed both in RSA analyses and in trial-level estimates of representational expansion, demonstrating that successful performance was associated with increased separation of task-relevant representations rather than a generic increase in neural discriminability. In contrast, models based on stimulus salience or simple centre-versus-edge distinctions provided substantially weaker relationships to the data. Together, these findings suggest that successful retrieval is associated with an ordered representational geometry in which behaviourally relevant distinctions are selectively amplified.

The interactions between belief, attention, and behaviour suggest a more general principle: that belief-based representations of task structure are associated with systematic patterns of information sampling across multiple timescales and modalities. This idea is formalised in the Sampling Emergent Attention (SEA) model (Braunlich & Love, 2022), which proposes that attention emerges through belief-guided sampling of information sources to maximise expected utility. Consistent with SEA, participants increasingly focused on task-relevant features as model-derived beliefs stabilised. However, our findings extend this framework by showing that selective sampling varied with inferred latent context rather than fixed stimulus features. Trial-wise prediction errors were associated with subsequent changes in gaze behaviour and neural context representations, suggesting that learning-related belief updates influence how information is sampled and represented. Furthermore, task-relevant information was selectively amplified within neural population activity, indicating that belief-guided sampling is associated with corresponding changes in representational structure. These findings are broadly consistent with the SEA framework and suggest that it may be extended to incorporate inference over latent structure.

While previous work has demonstrated that task-relevant features become increasingly emphasised in neural representations during categorisation (Duan et al., 2024; Mack et al., 2020), our study advances this literature in several important ways. Prior approaches often attribute representational compression to explicit task demands or data-driven feature selection. In contrast, we show that representational transformations covary with model-derived beliefs about latent structure rather than stimulus visibility or feature informativeness. Compared with Mack et al. (2020), which tracked dimensionality compression across learning blocks, we observed trial-by-trial changes in representational structure aligned with inferred context. Unlike Duan et al. (2024), which manipulated feature visibility, we held stimuli constant and instead manipulated latent task structure, allowing us to dissociate relevance from visibility. Relatedly, Flesch et al. (2022) demonstrated that extensive training leads to orthogonal context-specific manifolds in frontoparietal cortex. Our findings suggest that comparable transformations can emerge rapidly and reversibly as a function of trial-by-trial belief dynamics.

Together, these findings support an account of abstraction in which internal models, attention, and neural representations are closely coordinated during flexible behaviour. Rather than encoding all available information equally, participants appeared to infer a task-relevant subspace and selectively amplify distinctions within that subspace. In this view, latent-state inference does not simply reduce the dimensionality of the problem space; it restructures representational geometry such that behaviourally relevant dimensions become more separable while irrelevant dimensions are deemphasised. Such belief-guided representational reorganisation may provide an efficient mechanism for learning and decision-making in high-dimensional, non-stationary environments (Bellmund et al., 2018; Gershman & Niv, 2010; Konidaris, 2019; Leong et al., 2017; Mack et al., 2020; Niv, 2019; Radulescu et al., 2019; Schuck et al., 2016).

An open question is whether attention reflects the outcome of inference or whether it also contributes to learning-related changes in representation. Our findings are compatible with both interpretations. On one hand, attention closely tracked belief dynamics, consistent with a downstream correlate of model-derived inference. On the other hand, increased focus on relevant dimensions may itself contribute to greater differentiation of those dimensions in neural activity, consistent with a reciprocal relationship between attention and representational learning (Kruschke, 1996; Love et al., 2004; Nosofsky, 1986). This view resonates with recent work showing that hippocampal representations disentangle latent causes from stimulus identity to support generalisation (Courellis et al., 2024). Here, we extend this perspective by demonstrating coordinated relationships between model-derived belief dynamics, attentional selection, and dynamic changes in neural representational geometry (Flesch et al., 2022).

In sum, we propose that latent-state inference provides a useful framework for understanding flexible decision-making, being closely associated with both attentional allocation and neural representational structure. Across behaviour, gaze, and neural activity, participants selectively prioritised information relevant to the currently inferred context. As neural representations became aligned with inferred relevance, representational geometry was reorganised to selectively amplify behaviourally relevant distinctions, supporting abstraction, adaptability, and efficient control in complex environments (Bellmund et al., 2018; Gershman & Niv, 2010; Schuck et al., 2016).

## Acknowledgments

The authors would like to thank Burkhard Mess and Yvonne Wolff-Rosier for help in collecting data and customise scripts for optimal data collection and Max Schulz for translation of instructions. They would also like to thank Daniel Carlström Schad, Evie Coxon, Nicholas Menghi, Daniel Reznik and Simone Viganó for their constructive input and discussions on this work. This work was supported by the Max Planck Society (C.K. and C.F.D.) and the Minerva Fellowship of the Max Planck Society (S.T.)

## Data and code availability

All code used to conduct the analyses, the data and scripts to reproduce the main figures in the manuscript is available at Github (https://github.com/kerrencasper/Hidden-state-inference.git) and Zenodo (10.5281/zenodo.20537161).

## Materials and methods

### Subjects

Fifty participants (23 females, 27 males; mean age = 27.9 years, SD = 4.36 years, range = 19-36) took part in the study in exchange for financial compensation. All participants had normal or corrected-to-normal vision and reported no history of neurological or psychiatric disorders.

All experimental procedures, including behavioural piloting, were approved by the Ethics Committee of the Faculty of Medicine at Leipzig University, Germany (approval number: 211/23-EK), and were conducted in accordance with the Declaration of Helsinki. For the online piloting phase, participants provided informed consent by electronically ticking a consent box. For the MEG experiment, written informed consent was obtained prior to participation.

### Stimuli

The stimulus set consisted of 125 images of a car, each varying along three visual features (hereafter referred to as dimensions). These dimensions were: (i) the height of the rear wing, (ii) the distance between the car body and the wheels, and (iii) the brightness of the windows. These visual features were designed as perceptual proxies for abstract car attributes: wing height represented speed, wheel-body distance represented weight, and window brightness represented cost. Each dimension varied across five levels, resulting in a fully crossed set of 125 unique stimuli (5 x 5 x 5). Stimulus selection was partly motivated by prior research on geometric representations of similarity spaces (Gärdenfors, 2000), allowing us to manipulate dimensionality while focusing on goal-directed learning processes.

To ensure that perceptual spacing across levels was uniform within each dimension, we conducted a behavioural study employing maximum likelihood difference scaling (MLDS). This method estimates latent perceptual scale values based on participants’ dominance judgments across triads of stimuli (e.g., judgments of whether stimulus b is more similar to a or c). Participants were presented with a sufficiently large number of triads, and the resulting data were fit using maximum likelihood estimation (Supplementary Fig. 1).

This scaling task was piloted online using Pavlovia (via Prolific), with the experiment implemented in PsychoPy. Thirty participants took part (mean age = 29.23 years, SD = 4.77 years; 14 females). A repeated-measures ANOVA revealed no significant effect of relevant dimension on accuracy, F(2,58) = 1.86, p = .17, indicating that participants performed similarly across the weight, speed, and price dimensions. An unstandardised profile confirmed that the minimum and maximum scale values were comparable across dimensions, suggesting similar levels of difficulty and perceptual span for the three features. Crucially, this ensured that an equidistant cube, where each dimension spans an equal range from minimum to maximum, could reliably be used in the main task.

To enable potential future analysis of grid-like coding, we restricted stimulus pairs to specific spatial trajectories. These included cardinal angles (0°, 90°, 180°, 270°), oblique angles (45°, 135°, 225°, 315°), and symmetrical directions offset from cardinal axes by 20° and 60°, yielding a total of 16 angular trajectories. While the current study did not directly investigate grid-like representations, this design enables future analysis.

### Task

Prior to each phase of the experiment, participants received detailed instructions followed by training to ensure full understanding of the task requirements. The experiment comprised five phases: (1) a localiser task; (2) learning of the three stimulus dimensions; (3) learning the contexts’ reward-contingency preferences; (4) the serial reversal learning task; and (5) a post-task localiser. The primary focus of the current study was Phase 4 (the serial reversal learning task) in which participants were required to infer the current context and select actions accordingly.

### Localiser task

In the localiser task, participants were presented with individual images of cars and were instructed to press the space bar on the keyboard whenever the currently displayed car was identical to the one shown on the previous trial. These repeat trials, referred to as catch trials, served as an attention check and occurred approximately every 10 trials. Each trial began with a fixation cross (500 ms), followed by the presentation of a car image (1500 ms). All 125 unique car images were shown once, and 14 of them were repeated, resulting in a total of 139 trials. The localiser phase lasted approximately 6 minutes, including instructions and excluding breaks.

### Learning of different dimensions

In the second phase of the experiment, participants learned to discriminate within the different dimensions (i.e., visual features) of the car stimuli. They were informed that each dimension was a proxy for a hidden characteristic of the car (see Stimuli), but were not told how many levels each dimension contained. The goal of this phase was to implicitly learn the ordinal structure within each dimension through repeated exposure and feedback.

Before the first trial of each dimension block and then every 20th trial thereafter participants received a brief instruction screen indicating which feature to attend to (e.g., “Your task is to choose the fastest of the two cars!”, which referred to the rear wing). These instructions were displayed for 2000 ms. Each trial involved a two-alternative forced choice between two cars. Participants responded using the left and right arrow keys to select the first or second car, respectively. Trials followed the following structure: a fixation cross (750 ms), the first car (1500 ms), a second fixation cross (500 ms), and the second car, which remained onscreen until a response was made or for a maximum of 2000 ms. Every 20th trial, participants received feedback on their performance in the form of accuracy and the percentage of completed trials (presented for 2000 ms).

Participants completed 100 trials for each dimension, resulting in a total of 300 trials. If a participant achieved more than 80% accuracy and completed more than 80% of the trials within a dimension, they progressed to the next dimension. This phase lasted approximately 20 minutes, including two short breaks, assuming full completion of all 300 trials. Average performance was 97.22% (SE = .32%).

### Learning context preferences

In the third phase of the experiment, participants learned the car preferences of three fictional car-dealer characters, each representing a distinct context. The reward preferences were counterbalanced across participants to ensure that each dimension was relevant in isolation and in combination with another dimension across the three contexts. Specifically, for a given participant, one context exhibited preferences based on a single dimension (1D), while the other two contexts relied on combinations of two dimensions (2Da and 2Db). The relevant dimensions were partially overlapping across contexts. For example, if Context 1 rewards high speed (dimension 1), another context might reward low speed on the same dimension, resulting in anti-correlated reward contingencies. Alternatively, if the second context rewards a distinct combination like low speed and high cost (dimensions 1 and 3), this could reduce representational overlap and even approximate orthogonality in the reward structure.

Participants were informed, before the first and then every 20th trial, which dimension(s) were relevant for making correct decisions (e.g., “Your task is to choose the fastest and most expensive car!”). These instructions were presented for 2000 ms.

Each trial followed this structure: a fixation cross (750 ms), presentation of the first car (2500 ms), a second fixation cross (500 ms), and the second car, which remained onscreen until a response was made. Participants responded using the left and right arrow keys to select the first or second car, respectively. Feedback was given after each trial for 1500 ms if the participant made the correct choice. Unbeknownst to participants, feedback was determined by the underlying stimulus-reward contingency maps specific to each context (see Fig. 1). Thus, the goal was to implicitly learn the reward structure associated with each context’s preferences. After every 20th trial, participants received feedback on the percentage of the current phase completed.

In total, participants completed up to 375 trials (125 per context). However, if they reached a performance threshold of over 85% accuracy and completed at least 70% of the trials (i.e., 88 trials), the current context phase ended early, and after a short break, training resumed with a new context. This adaptive thresholding procedure limited the total duration of this phase to approximately 30-40 minutes, including two breaks. Average performance was 91.32% (SE = .51%).

### Serial reversal-learning task

In the fourth and main phase of the experiment, participants were tested on the knowledge acquired during the previous phases. On each trial, participants were required to make a decision without knowing in advance which context they were currently playing. Instead, they had to infer the correct context through trial and error based on the feedback received after each decision. Unbeknownst to participants, they remained in the same context for between 5 and 8 trials before switching to another. These block lengths were chosen to maximise the number of trials and context switches while keeping the overall task duration within reasonable limits to minimise participant fatigue. Each trial began with a fixation cross of random duration (1000, 1250, or 1500 ms), followed by presentation of the first car image (2000 ms), a second fixation cross (500 ms), and then the second car, which remained onscreen until a response was made. Participants responded using the left or right arrow key to choose the first or second car, respectively. Feedback was provided for 1500 ms if the choice was correct.

Due to confusion and low behavioural performance in a separate piloting task, we added instructions when a context was about to change. Accordingly, when a context switch was about to occur, participants were shown an instruction screen reading: “There is now a high probability that there will be a shift in the character you are playing!” (German: “Es besteht nun eine hohe Wahrscheinlichkeit, dass sich der Charakter, den du spielst, verändert!”). This warning served two purposes: (i) it ensured that the first trial of a new context was behaviourally equivalent across switches (i.e., 50% chance accuracy), and (ii) it allowed us to control for dimensionality and behavioural uncertainty in our subsequent analyses (see Analysis section for details).

After approximately every 30 trials (timed to avoid disrupting context switches), participants received information about the percentage of the phase completed. In total, participants completed 351 trials, including 54 context switches. This phase lasted approximately 40 minutes, including short breaks.

## Experimental setup

### Data acquisition

The piloting task was implemented online using custom-written code in Python and JavaScript in PsychoPy, and tested via Pavlovia (https://pavlovia.org). Participants were recruited through the online platform Prolific (https://www.prolific.com). The main experiment was implemented using custom-written code in MATLAB (MathWorks), with PsychToolbox (Brainard, 1997) and the EyeLink Toolbox (Cornelissen et al., 2002) for stimulus presentation and eye-tracking integration.

In the main task, stimuli were presented relative to the monitor’s refresh rate to ensure precise timing. During the main experiment, participants were seated in front of a monitor with a resolution of 1280 x 1024 pixels at a viewing distance of approximately 90 cm, with their heads positioned within the MEG helmet to minimise head movements. Eye movements were recorded continuously using an EyeLink 1000 Plus system (SR Research) at a sampling rate of 1000 Hz. The eye tracker was positioned 58 cm from the participants’ eyes. Before each experimental phase, the eye tracker was calibrated using the standard 9-point calibration and validation procedures provided by the EyeLink software.

### Analysis

The general hypotheses and analysis plan were pre-registered prior to data collection and are available on the Open Science Framework (OSF; https://osf.io/6×42q/?view_only=4e3c9567d6a84ea2863e609677c76627). Data preprocessing was conducted using the MNE-Python toolbox (Gramfort et al., 2013) and custom-written scripts in Python. The main time-resolved neural analyses were carried out using the FieldTrip toolbox (Oostenveld et al., 2011) and custom-written MATLAB code. A total of 50 participants took part in the MEG experiment. For MEG analyses, no participants were excluded due to excessive artefacts or poor data quality, resulting in a final sample of 50 participants included in the reported analyses.

### Statistical analysis

In accordance with the pre-registration, all main statistical tests that were mentioned in the pre-registration were one-sided and conducted at the group level. Non-parametric cluster-based permutation tests were used to assess significance when applicable.

### MEG data analysis

MEG data were recorded at the Max Planck Institute for Human Cognitive and Brain Sciences (MPI CBS) in Leipzig, Germany, using an Elekta Neuromag TRIUX system (Elekta, Stockholm, Sweden) comprising 306 channels; 204 planar gradiometers and 102 magnetometers, sampled at 1000 Hz.

### Oculomotor analysis

Oculomotor behaviour was converted from edf to MATLAB using the Edf2Mat toolbox (https://github.com/uzh/edf-converter) and downsampled to 250Hz, to conform to the same sampling rate as the MEG signal. We interpolated time points where the eye-tracking signal was not available (e.g., due to blinks) and set time points where participants looked beyond the field of view of the eye-tracker, automatically detected by the Eyelink, to NaNs. We then set each time point after each trial’s reaction time to NaN. For all our oculomotor analyses, we constructed three regions of interest (ROIs; the wheels, the windows, and the rear wing; see Supplementary Fig. 2), in which we assessed gaze and fixation.

To quantify selective attention dynamics across learning, we examined the proportion of gaze time allocated to attended versus unattended feature dimensions over the course of trials within each context block. We analysed gaze behaviour separately for 1D and 2D trials, and then combined the results (Fig. 4A).

For each subject, gaze time was extracted from the late phase of the trial (2.5-4 seconds post-stimulus onset). Gaze samples were binarised within predefined regions of interest (ROIs) corresponding to the attended and unattended dimensions, based on the reward structure of each trial. To account for overlapping features across context pairs (e.g., 1 and 3 and 2 and 3), fixation within an unattended ROI was only counted when it reflected unique information not included in the attended dimensions. Similar pattern as in 3A was observed when instead using fixations within ROI (early: M = 1.28 dimensions, SE = .02; late: M = 1.17 dimensions, SE = .01; t(49) = 7.99, p < .01)

Gaze proportions were computed for each trial position (1-8) within a context block. For each trial, we calculated the total number of gaze samples falling within attended ROIs and separately for each of the two unattended dimensions. We then computed the proportion of total gaze time attributed to attended and unattended dimensions, averaging across all valid trials at each position.

This was done separately for 1D and 2D trials, and the resulting matrices of attended and unattended gaze proportions were concatenated across subjects. For visualisation, we computed the mean and standard error of the mean (SE) across participants, and plotted the resulting time courses (Fig. 4B).

The same analysis as in 3B was conducted but now splitting the data into correct and incorrect trials, and calculate the gaze proportion within relevant dimensions (Fig. 4C).

Inference trials were calculated by contrasting trials in which the made a correct choice after an initial error or whether they continued to be incorrect. We used a time-resolved analysis and corrected for multiple comparisons using a cluster-based permutation test (Fig. 4D).

Our main pre-registered hypothesis was to assess the entropy across trials within a context (Fig. 4E). To assess this, we counted the probability of ones and zeros (present/absent) within the ROIs and estimated entropy for correct and incorrect trials separately and for each trial within a context using the formula:

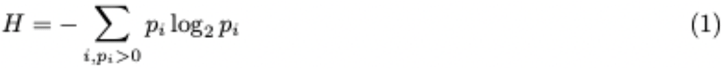

where *H* represents entropy, p_i_ is the probability of gaze presence in an ROI. The summation runs over all nonzero probabilities (p_i_ > 0), as probabilities of zero do not contribute to entropy.

Trial 1-5 within a context contained enough trials for both correct and incorrect responses to reliably estimate entropy.

### Preprocessing

For each participant, bad channels were annotated in a separate document, which was incorporated into the main data structure prior to preprocessing. To attenuate environmental noise and correct for head movements, we applied Signal Space Separation (SSS) using the Maxwell filter. The origin of the head position was set to auto with respect to the head coordinate frame. An internal expansion order of 11 and an external expansion order of 2 were used. Magnetic scaling was automatically determined by the software (mag scale = auto). Site-specific calibration and crosstalk correction files were applied throughout the filtering process.

Channels were automatically rejected if they exceeded specific amplitude thresholds: 400 fT for magnetometers 4000 fT/cm for gradiometers, and 150 µV for EOG channels. Channels identified as bad during preprocessing were excluded from the SSS computation to avoid introducing artefacts into the spatial filter. Head movement correction was performed using a sliding window algorithm with 200 ms windows and 10 ms steps. The HPI coil fit was constrained by an error threshold of 5 mm and a g-value of 0.98. Following movement correction, the MEG signals were virtually re-aligned to the participant’s mean head position during the initial localiser task to ensure comparability of sensor-level data across all experimental phases. After applying the MAX filter, all data were visually inspected to confirm that artefacts had been sufficiently attenuated and that no signal distortions had been introduced. The cleaned data were then band-pass filtered with a high-pass cut-off of 0.1 Hz and a low-pass cut-off of 200 Hz. Line noise was attenuated using notch filters at 50, 100, and 150 Hz.

Artefact rejection was further refined using independent component analysis (ICA). To optimise the decomposition, a copy of the data was high-pass filtered at 1 Hz and downsampled to 250 Hz. ICA was performed using the *fastica* algorithm from the MNE-Python toolbox, with the maximum number of components set to 100, a maximum of 300 iterations, and a fixed random state for reproducibility. Bad channels were excluded from ICA. Components reflecting eye movements and cardiac activity were manually identified and removed. The cleaned data were then back-projected to the original signal space.

Next, the continuous data were epoched into trials. We used the *AutoReject* algorithm (Jas et al., 2017) as implemented in MNE-Python to automatically detect and reject bad trials and repair noisy channels. Each participant’s preprocessed data was then visually inspected to confirm the quality of the trial rejection and channel interpolation. On average, 327.28 trials (SE = 2.99; range: 253-351) out of the total 351 trials were retained for the serial reversal learning phase.

Finally, the cleaned and epoched data were transferred to MATLAB and analysed using the FieldTrip toolbox for time-resolved and multivariate analyses.

### Context decoding

Data were preprocessed by first smoothing it using a 200 ms moving average (implemented via smoothdata in MATLAB), applying multivariate noise normalisation and cocktail blank removal to increase the signal-to-noise ratio, following methods established in previous multivariate work (Guggenmos et al., 2018). On each trial, the class label corresponded to the currently active latent context and coded as one of three possible contexts. Decoding was performed independently at each time point from -200 to 4000ms. At each time point, the feature vector consisted of activity across all selected MEG channels.

For each participant and time point, we trained a multiclass linear one-versus-all ridge-regularised logistic regression classifier, implemented in MATLAB. Classification performance was estimated using 3-fold cross-validation and repeated five times. Within each fold, training and test data were were z-scored. Decoding accuracy was defined as the mean of the class-wise accuracies across the three contexts. Accuracy estimates were then averaged across repetitions.

In addition to decoding accuracy, we extracted trial-wise neural evidence for the true context. For each held-out trial, we obtained the classifier score assigned to the trial’s true context label. This produced a time-resolved measure of neural evidence for the currently active latent context on each trial. These true-context evidence values were subsequently used in trial-wise regression analyses relating neural context representations to model-derived belief dynamics.

To assess whether neural context evidence was modulated by computational variables from the hidden-state inference model, we fit linear regression models at each time point and participant. Neural evidence for the true context was predicted from model-derived belief strength, entropy, and unsigned prediction error. In a second model, we tested whether context evidence on the current trial was influenced by prediction error on the previous trial, by replacing current prediction error with PE t-1. These models were also fit separately for correct and incorrect trials, allowing us to test whether the relationship between prior prediction error and current neural context evidence differed as a function of behavioural outcome.

### Representational similarity analysis

Data were preprocessed by first smoothing it using a 200 ms moving average (implemented via smoothdata in MATLAB), applying multivariate noise normalisation and cocktail blank removal to increase the signal-to-noise ratio, following methods established in previous multivariate work (Guggenmos et al., 2018). In line with the pre-registration, and to estimate spectral power, we convolved the time series with complex Morlet wavelets. Power was calculated using linearly spaced wavelets from 3 to 100 Hz: all integer frequencies between 3 and 30 Hz, and every fifth frequency between 40 and 100 Hz. The continuous-time wavelet transforms were squared and log-transformed. To normalise for power differences across frequency bands, we z-scored power values separately for each frequency using that frequency’s mean and standard deviation across all trials. These were then averaged within and across canonical frequency bands: theta (3-7 Hz), alpha/beta (8-30 Hz), and gamma (40-100 Hz), resulting in a single signal representing the average across all frequencies. We split the dataset based on task demands into trials where participants focused on one dimension (1D) and those where they focused on two dimensions (2D).

To characterise the structure and temporal evolution of neural representational geometry, we constructed a series of model representational dissimilarity matrices (RDMs; Fig. 6A; Supplemental Fig. 5 and 6). Five model geometries were generated for both 1D and 2D task contexts: linear, extreme-expanded, extreme-versus-middle, salience-U, and centre-versus-edge. Each model assigned distances between stimulus values according to a distinct hypothesis regarding the organisation of representational space. Pairwise dissimilarities between all stimuli were calculated as Euclidean distances in the corresponding model space and normalised to the maximum distance within each RDM.

Neural RDMs were computed from MEG activity using a sliding-window approach. For each trial, neural activity during an early period (250-500 ms after first-stimulus onset; time window extracted from significant context decoding in Figure 5B) was compared with activity during the later period (2500-4000 ms after first image onset; from 0 to 1500 ms after second-stimulus onset). Temporal response patterns were extracted within overlapping 200-ms windows (80% overlap), and pattern similarity between the early and late windows was quantified using Pearson correlation across time points. Neural dissimilarity was defined as one minus the correlation coefficient. This yielded a trial-wise neural distance measure for every sensor and time window.

To construct stimulus-level neural RDMs, trial-wise dissimilarities were averaged across all trials corresponding to each stimulus pair within the 5 x 5 x 5 stimulus space, producing a 125 x 125 neural RDM for each sensor and time window. Neural RDMs were first averaged across the encoding windows and then vectorised. The resulting neural RDMs were compared with each model RDM using Spearman rank correlation, yielding a time-resolved measure of correspondence between neural and model representational geometry.

To quantify representational expansion directly, we additionally estimated a trial-wise expansion coefficient. For each participant, sensor, and time window, neural distances were regressed against distances in the task-relevant stimulus space. For 1D analyses, distances were computed along the currently relevant stimulus dimension; for 2D analyses, distances were computed within the relevant two-dimensional stimulus space using Euclidean distance. The resulting regression slope provided a measure of how strongly neural representational distances scaled with distances in the behaviourally relevant stimulus space, with larger positive values indicating greater separation of task-relevant representations.

For statistical analysis, neural-model correlations and expansion coefficients were compared between correct and incorrect trials using non-parametric cluster-based permutation testing across sensors and time. Separate tests were performed for each of the 5 model RDMs (corrected p-value = .01). To assess whether observed neural-model correlations exceeded those expected by chance, we additionally generated participant-specific null distributions by randomly permuting the rows and columns of the model RDM while preserving its symmetry and overall distance distribution. This procedure was repeated 250 times per participant, and empirical effects were evaluated relative to the resulting null distributions.

## Computational modelling

### Model Description

We evaluated four computational models to account for participants’ behaviour in the serial reversal learning task. Three of these models were variants of Q-learning, a family of reinforcement learning (RL) models based on value updating. The fourth model was a hidden state inference (HSI) model, which has recently shown promise in a similar task with rodents. The modelling objective was to predict participants’ inferences about the current context and their button-press decisions.

### Q-learning

We implemented a context-sensitive Q-learning model that simultaneously tracked beliefs about context identity and action values. On each trial, the model maintained context-specific Q-values and used softmax inference to estimate the current context. Let ct ∈{1,2,3} denote the latent context on trial t. Q-values were initialised uniformly:

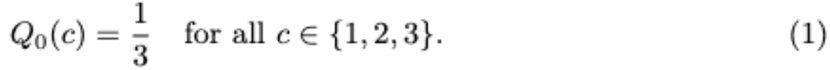

The probability of inferring context c was computed via a softmax decision rule:

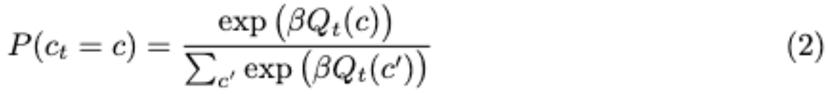

After observing the true context and reward on each trial, the model updated the Q-value for the true context using a reward prediction error

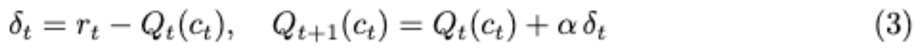

### Q-learning with Forgetting

A second model variant extended standard Q-learning by incorporating forgetting, allowing the model to gradually reset the value of non-chosen contexts toward the average expected value. This mechanism accounts for the idea that unused associations decay over time unless reinforced. After updating the Q-value for the true context using the standard Q-learning rule:

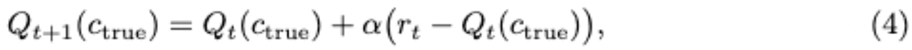

the model adjusted the Q-values of the non-chosen contexts based on their deviation from the average value Qt. Specifically, for each non-chosen context c’ ≠ c_true_, the Q-value was updated according to

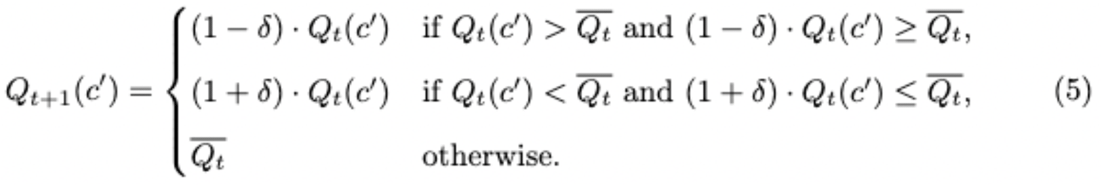

### Q-learning with Counterfactual Updating

A third model extended standard Q-learning by incorporating counterfactual updating. In this model, not only is the value of the chosen context updated based on received feedback, but the values of the unchosen contexts are also adjusted in the opposite direction. This mechanism captures the idea that selecting one context and receiving a particular outcome can inform beliefs about the unselected alternatives. For the chosen context c_true_, the update rule depended on the received reward r_t_:

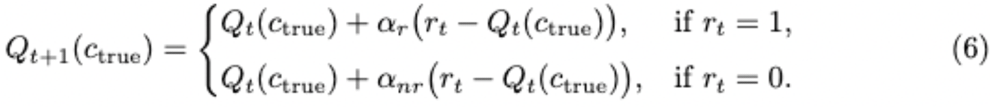

For each unchosen context c’ ≠ c_true_, the update was based on the counterfactual outcome 1 - r_t_, again with separate learning rates for rewarded and unrewarded trials:

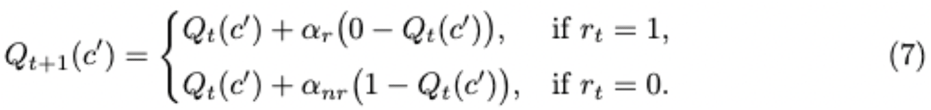

### Hidden State Inference Model

The Hidden State Inference (HSI) model adopts a Bayesian framework, in which agents infer the most likely latent state (i.e., context) of the environment on each trial. The agent maintains a belief distribution *b*^*t*^*(sJ = p(s*_*t*_ |*o*^*t-1*^) over possible states s ∈ {1,2,3}, conditioned on past observations o^t-1^. Each observation is defined as an action-reward pair:

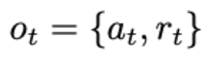

On each trial, beliefs are updated via Bayes’ rule, combining the transition model and the observation likelihood:

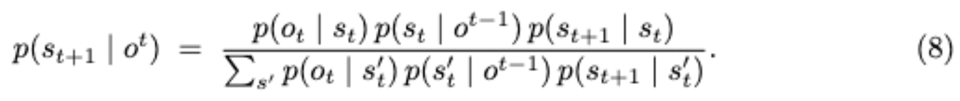

State transitions follow a Markov process with persistence parameter *γ* ∈ [0, 1]. The probability of remaining in the same state is

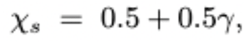

and the remaining probability 1-χ _s_ = 0.5-0.5 γ is split evenly across the two alternative states, yielding

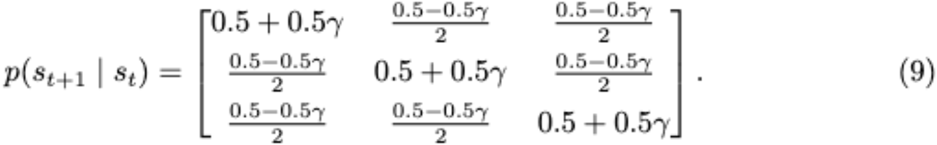

Let a^*^(*s*) denote the action that is rewarded in state *s* (the state’s “correct” action in the task). We introduce two sensitivity parameters *c, d* ∈ [0, 1] controlling how strongly the likelihood favours feedback that is consistent or inconsistent with the current state’s contingency. The likelihood *p*(o_t_ | *s*) depends on whether the taken action matches a^*^(*s*) and on the received reward r_t_:

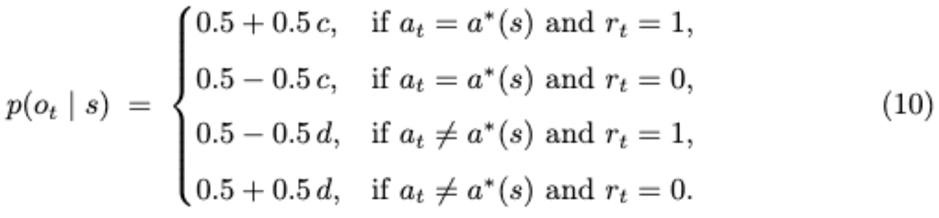

Note that for each model on each trial, we renormalised the context values so they represent relative beliefs, by dividing by the sum of all values.

### Action Selection Based on Context Inference

Although each model used a different mechanism to infer the current context, via value updating, decay, counterfactual reasoning, or Bayesian inference, the final step of action selection was shared across all models. On each trial, once the model computed a probability distribution over contexts *P(c*_*t*_) or beliefs over latent states *b*_*t*_*(s*), it used these to estimate the expected reward for each action. Specifically, expected reward values for the left and right choices were computed as the weighted sum of the rewards in each context:

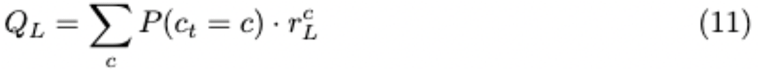

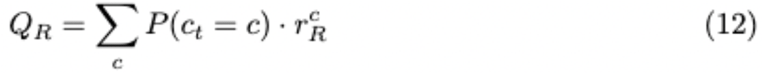

Here, 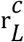 and 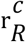 denote the reward associated with the left and right choices, respectively, in context *c* (or latent state *s* in the case of the HSI model).

The probability of selecting each action was then determined using a separate softmax function:

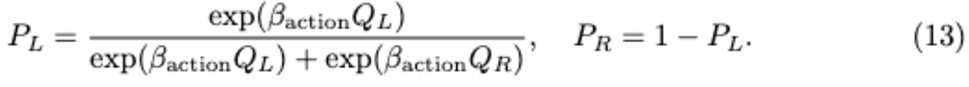

The model then simulated a button press based on these action probabilities. These predicted responses were used to compute trial-by-trial likelihoods and to evaluate how well each model captured participants’ observed behaviour. Thus, the model was more confident in selecting the correct action when the inferred probability for the corresponding context was high, as this increased the weighting of the relevant reward contingency in the expected value computation.

### Model Fitting

To estimate the parameter values that best characterise the behavioural data, we employed a likelihood maximisation approach to model fitting. For each behavioural session, we estimated the probability of individual choices based on a given model (*m*), its parameters (Θ _m_), and the history of choices and outcomes within that session.

The logarithms of the choice probabilities corresponding to the participants’ choices on each trial were then summed. The MATLAB function *fmincon* was used to determine the parameter values that minimised the negative log-likelihood of the data (*LL*) given the model parameters:

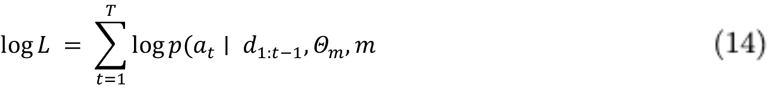

To prevent convergence to local minima in the minimisation process, model fitting was repeated 50 times using randomly selected initial values within predefined bounds for each parameter. The log-likelihood of the best-fitting model was recorded for each run, and the final parameter estimates were taken from the run with the highest log-likelihood value.

To identify the model that provided the most parsimonious fit to the data, we compared the model fits for each individual session using the Bayesian Information Criterion (BIC). The BIC incorporates an explicit penalty for the number of free parameters (k_m_) in the model m, thereby controlling for overfitting:

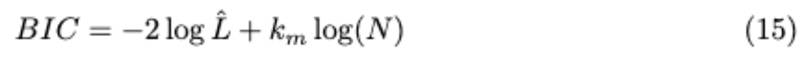

where *L* denotes the log-likelihood value at the best-fitting parameters, and *N* represents the number of trials within a session. To compare model fits for each participant, we calculated the difference in BIC scores (*LBIC*) between each model fit and the most parsimonious model for the same session.

### Model verification

To assess how reliably the best-fitting model from our model comparison procedure reflects the true generative process, we performed a model recovery analysis (Wilson & Collins, 2019). This involved a comprehensive all-by-all comparison: for each model, we simulated behavioural data and then fit all models to each simulated dataset.

We quantified the proportion of datasets generated by one model that were best fit by each of the candidate models (i.e., fit model and simulated model), and summarised the results in a confusion matrix. If each model perfectly recovered itself, the confusion matrix would be an identity matrix, indicating perfect recoverability.

We also computed the corresponding inversion matrix, which reflects the probability that a given model generated the data, given that it provided the best fit (i.e., simulated model and fit model). This allows us to estimate how confidently we can infer participants’ underlying strategies based on model fits and specifically, how likely it is that the best-fitting model also generated the observed behaviour.

### Alignment with pre-registration

The pre-registration was done June 19, 2023. The two first main hypotheses were tested in the way described in the pre-registration: For behaviour we measured accuracy and reaction times (RTs) and used computational modelling. We pre-registered to use linear regression models to measure an increase/decrease in accuracy/RTs. For oculomotor behaviour we used entropy, gaze, fixations and heat maps, but did not use scan paths. For MEG, we used the features that we pre-registered (decomposed data with all frequencies averaged) to measure the similarity between model representational dissimilarity matrices (RDMs) and brain data. We created 1D and 2D RDMs to assess expansion and compression. However, due to low trial count, we were not able to assess the dimensionality in the first trial of a new context using a 3D matrix. We were also not able to test the third main hypothesis, due to the experiment already being very long and cognitively demanding for participants. Therefore, the last part about generalisation had to be taken out.

We decided against using source-reconstruction and ROIs. This was mainly due to the paper already so dense and conducting these analyses would have substantially increased the length.

Although we intended to use principal component analysis (PCA) as a measure of neural compression, we implemented the equivalent procedure using singular value decomposition (SVD). SVD and PCA are mathematically equivalent methods for low-rank approximation: PCA decomposes the covariance matrix of the data, whereas SVD directly decomposes the data matrix itself. We adopted the SVD formulation because it allows condition-mean activity patterns to be decomposed and cross-validated, as recommended by Ahlheim and Love (2018) and Liang et al. (2025). This cross-validated SVD procedure yields the same latent subspace as PCA but provides a more interpretable and numerically stable estimate of representational dimensionality. Thus, the analysis remains consistent with the preregistered aim of quantifying neural compression, differing only in the specific but equivalent mathematical implementation.

Since the pre-registration, we have accumulated new evidence regarding our analyses. First, the seminal paper by Mishchanchuk et al. (2024) showed that in a similar task, mice’s behaviour was best captured by a Hidden state inference model. Therefore, we added this to the suggested reinforcement learning models. Secondly, the modelling part became much more extensive due to the surprisingly good alignment between the model and data. We therefore decided to further investigate model-derived vaiables in oculomotor behaviour and neural data.

Although an increase in frontal theta power was hypothesised for early vs. late trials within a context in the pre-registration, we decided to continue on the laid out path from the modelling and oculomotor analyses, instead contrasting first trial vs rest within a context, as we understood this to be a very strong predictor of cognitive control demands.

## Supplemental analyses

### Frontal midline theta

To provide an independent neural validation of the latent-state inference signals captured by the HSI model, we examined frontal midline theta (FMT) activity, which has been implicated in uncertainty monitoring, cognitive control, and behavioural adaptation (Cavanagh & Frank, 2014; Cavanagh & Shackman, 2015). We performed time-frequency decomposition on MEG signals using complex Morlet wavelets. Preprocessed sensor-level data were epoched from -1000 to 5000ms and analysed separately per trial. We focused on frontocentral gradiometers (all gradiometers anterior of midline). For each trial, we estimated spectral power from 2 to 20 Hz in 1-Hz steps, using variable-width wavelets with a minimum of three cycles and ∼500 ms window length. To reduce inter-trial and inter-frequency variability, we baseline-corrected power using a pre-stimulus window (-500 to -100ms), expressed as relative change. Theta-band power (3-7 Hz) was squared and log-transformed and then averaged across frequency bins, and z-scored across time within trial. This yielded a time-resolved measure of FMT amplitude per trial and sensor. Next, we grouped the trials into 1) first trials of a context and 2) all other trials and performed a non-parametric cluster-based permutation. Stimulus-locked analyses revealed significantly greater midfrontal theta power on the first trial compared to subsequent trials within a block (p_clusters_ <.05; Supplemental Fig. 3A), consistent with increased uncertainty and the need to update latent task representations following a context change.

### Frontal midline theta and prediction error

To examine trial-wise modulations in frontal theta activity, we processed the data as in previous section, but instead epoched trials from -1000 to 5000ms around outcome onset and analysed separately per trial. We again focused on frontocentral gradiometers (all gradiometers anterior of midline), based on prior work implicating frontal midline theta (FMT) in feedback and outcome processing (Cavanagh & Frank, 2014); estimated spectral power from 2 to 20 Hz in 1-Hz steps, using variable-width wavelets with a minimum of three cycles and ∼500 ms window length; and baseline-corrected power using a pre-outcome window (2000 - 2400ms), expressed as relative change, in line with (Lopez-Gamundi et al., 2024). Theta-band power (3-7 Hz) was squared and log-transformed and then averaged across frequency bins, and z-scored across time within trial. This yielded a time-resolved measure of FMT amplitude per trial and sensor, suitable for regression against model-derived prediction error.

To test whether frontal theta activity varied as a function of model-derived signed PE, we regressed single-trial frontal theta amplitude against trial-wise signed PE estimates from the HSI model. Trials were aligned to response onset, and time windows spanning from 200 ms before to 1000ms after response were extracted per trial. For each participant, we performed linear regression at each sensor and time point, using signed PE as the predictor and post-response theta amplitude as the dependent variable. The resulting beta maps reflected how strongly theta power tracked the magnitude and direction of the PE. To identify significant spatiotemporal clusters, we used a cluster-based permutation test as implemented in the Fieldtrip toolbox. This analysis revealed a significant frontal cluster, indicating that theta activity systematically covaried with model-derived signed PE (Supplemental Fig. 5B).

To further probe this effect, we grouped trials into percentiles (0-33, 34-66, 67-100) based on signed PE (negative, neutral, positive) and averaged theta power over the significant cluster. Theta power was significantly greater following positive than negative PEs (positive: M = .04, SE = .003; negative: M = .02, SE = .004; t(49) = 3.39, p = .0014), indicating that frontal theta was sensitive not only to the magnitude of surprise but also to outcome valence (Supplemental Fig. 3C).

Finally, we asked whether individual differences in PE-related theta signalling were associated with behavioural performance. Across participants, stronger coupling between signed PE and frontal theta predicted higher task accuracy (r = .34, p = .02), suggesting that participants whose frontal theta more strongly tracked model-derived outcome signals performed better overall (Supplemental Fig. 3D).

### Singular value decomposition

We estimated the effective dimensionality of neural activity patterns using a cross-validated singular value decomposition (CV-SVD) procedure adapted from Ahlheim and Love (2018) and Liang et al. (2025). For each participant, trial x sensor activity patterns were extracted separately for an early (0 to 2000ms) and late (2500-4000ms) time window (as there were different stimuli shown in the two separate time windows). Trials were labelled according to the discrete stimulus levels on each dimension (1-5), the attended dimensions on that trial, and the model family being evaluated (i.e., 1D, 2D and per-axis 1D). Labels were assessed on a per-model basis, and any conditions with fewer than three repetitions were excluded. For each participant and model, the resulting trial x sensor matrices were partitioned into K = 3 cross-validation folds using a label-stratified, run-aware procedure that ensured (i) each fold contained examples of each stimulus class whenever possible, and (ii) trials from the same run (one context) were kept within the same fold. In each iteration, K - 1 folds served as the fitting set and the remaining fold as the validation set. Condition-mean activity patterns were computed separately for the fitting and validation data and aligned using only the set of conditions present in both folds. Before decomposition, both fitting and validation matrices were z-scored using the mean and standard deviation estimated from the fitting set only, ensuring strict train-test separation.

We then decomposed the fitting condition-mean matrix into orthogonal components using SVD. The fitted data were reconstructed at increasing dimensionalities (from 1 up to a model-specific maximum determined by the number of conditions, sensors, and singular values), and model performance was quantified as the Pearson correlation between the reconstructed fitting data and the unscaled validation data. For each fold, the dimensionality estimate was defined as the number of components that maximised this cross-validated correlation. This entire procedure was repeated 50 times with new fold assignments, yielding distributions of best-k values, cross-validated reconstruction curves, and corresponding split-half reliability estimates for each model and time window. The resulting metrics were averaged across all folds and repeats to obtain a robust cross-validated estimate of representational dimensionality and reconstruction quality. This measure indexes how well a low-dimensional representation of the condition-mean activity patterns generalises to unseen data within each participant (Supplementary Fig. 4A).

To assess the relationship between representational dimensionality and stimulus separability, we performed decoding analyses on the same trial x sensor matrices used for SVD, using exactly the same fold assignments to preserve comparability. Decoding was performed using a multinomial logistic regression classifier with an L2 (ridge) penalty, implemented in MATLAB. As in the SVD analysis, training and test features were z-scored using parameters estimated from the training data only. Because some stimulus classes were absent in certain test folds, particularly for two-dimensional or per-axis models, we computed balanced accuracy only over the classes present in the test fold, avoiding artificial penalties for classes with no held-out examples. For each participant, decoding accuracy was averaged across repeats and folds and then related to their corresponding dimensionality estimates (Supplementary Fig. 4B-C).

Group-level significance for the dimensionality-decoding relationship was assessed using permutation tests in which participant labels were randomly shuffled 1000 times to generate a null distribution of correlations. Early and late time windows were combined by averaging participant-wise estimates across windows prior to computing group-level statistics.

To test hypotheses about the structure embedded in these neural manifolds, we evaluated four models that reflect the task’s conceptual geometry. An attended 1D model, comprising stimuli varying along a single attended dimension (five levels), tested in 1D trials; an attended 2D model, spanning all combinations of two task-defined dimensions (5 x 5 levels); and two separate 1D models for each of the attended axes in 2D trials. Significance was assessed using a within-subject permutation test that re-ran the full pipeline after shuffling stimulus labels within cross-validation folds (250 permutations). The resulting null distributions entered a second permutation level at the group stage (1000 permutations).

Correct 1D trials were well captured by the attended 1D model (mean reconstruction correlation = .07, SE = .01, z = 31.65, *p* < .01), with an average of 3.36 effective dimensions (SE = .48) (Supplementary Fig. 4A), whereas incorrect trials produced a reliably lower ED (41/50 participants included with enough incorrect trials; mean reconstruction correlation = .062, SE = .01, z = 13.85, *p* < .01, ED = 2.43, SE = .38; Welch’s unequal-variance t-test between correct and incorrect ED, t(65.53) = 6.23, p < .01).

For correct 2D trials, the joint 2D model provided a reliable fit (mean reconstruction correlation = .029, SE = .004, z = 5.33, *p* < .01), with an average of 10.32 ED (SE = 1.46). Incorrect trials again yielded lower ED (mean reconstruction correlation = .037, SE = .005, z = 3.92, *p* < .01, ED = 3.67, SE = .52; Welch’s unequal-variance t-test between correct and incorrect ED, t(59.82) = 9.98, p < .01). The two independent 1D axes also fit well (axis 1, mean reconstruction correlation = .07, SE = .01, z = 31.96, *p* < .01, ED = 3.26, SE = .46; axis 2, mean reconstruction correlation = .08, SE = .01, z = 37.22, *p* < .01, ED = 3.19, SE = .45). This pattern is consistent with a linearly separable representation along each attended dimension, embedded within a larger joint 2D manifold.

Finally, we tested whether higher ED supports better stimulus separability, motivated by the idea that higher dimensionality may support finer discrimination. ED was positively correlated with decoding accuracy for correct 1D trials (ρ=.21; exceeding the null distribution generated by 1000 permutations, z = 2.18, p = .025; Supplementary Fig. 4C), and for the independent axes of 2D trials (ρ = .29, permutation test: z = 2.86, p = .002; Supplementary Fig. 4C). No such relationship emerged for incorrect trials or for the joint 2D manifold (ρ = -.10, p = .31).

Together, these findings show that neural representational dimensionality varies systematically with task demands and behavioural performance. Higher dimensionality was observed in conditions requiring integration across multiple task-relevant features and on trials where participants responded correctly, suggesting that task-relevant information is more strongly expressed in neural population activity under these conditions. These results are consistent with the idea that neural representations are organised in a manner that reflects task-relevant structure.

## Supplementary material

**Figure S1.**
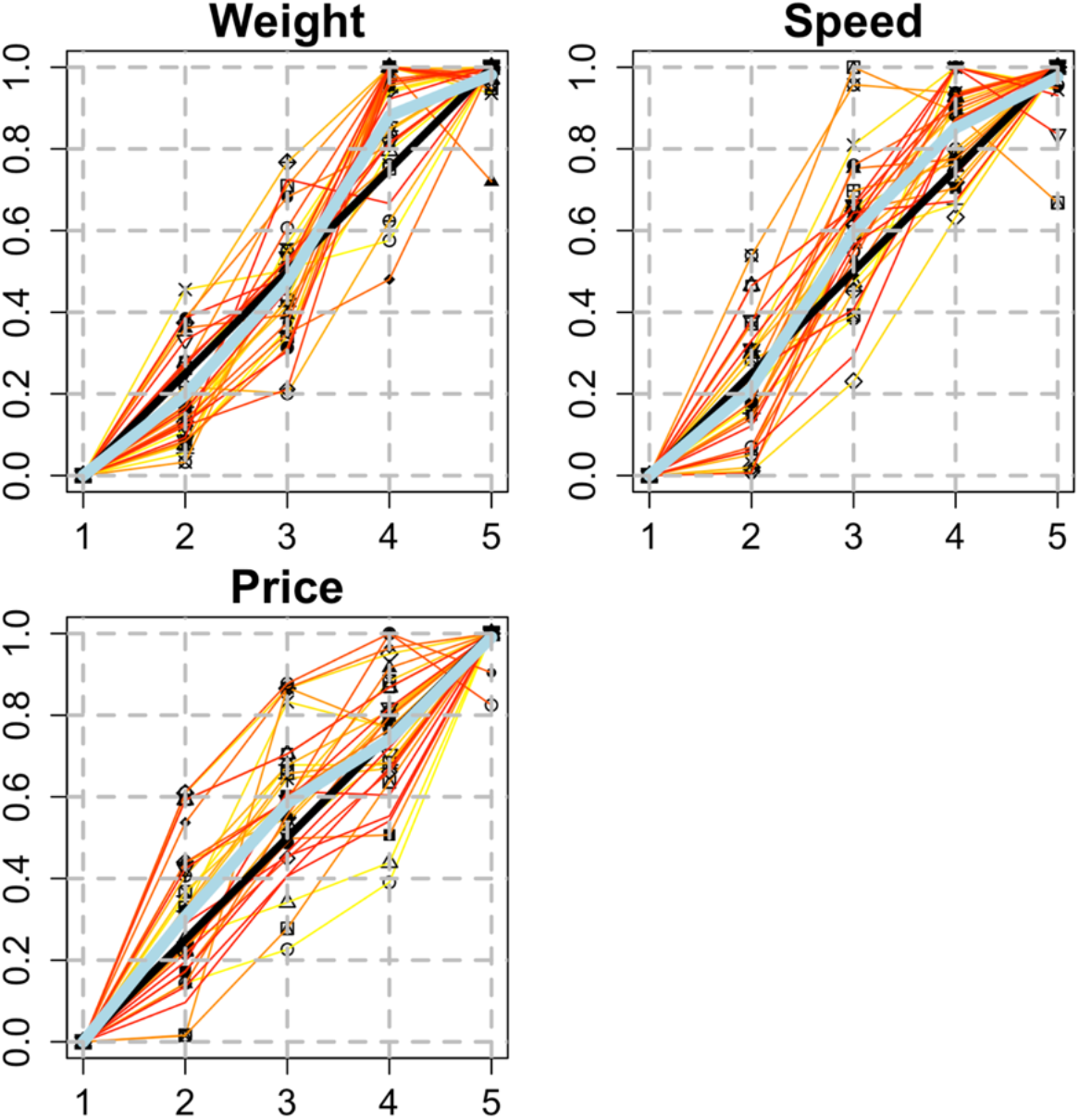
Maximum likelihood difference scaling of different dimensions. A pilot study comprising 30 participants assured that the psychological distance between levels within each dimension was equal. The light blue diagonal shows the average for all participants for any given dimension, whereas the black shows the optimal performance. Lines in different colours show each participant’s performance. If the line is above the diagonal, it means that the psychological distance was judged larger, whereas when the line was below the diagonal, the distance was judged to be smaller.

**Figure S2.**
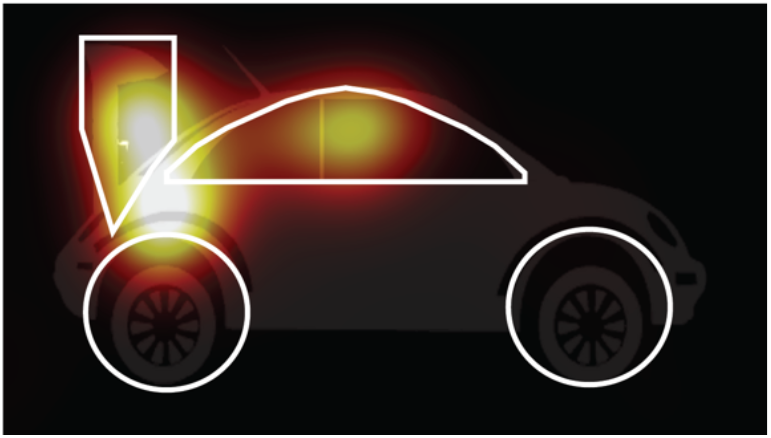
Regions of interest for oculomotor analyses and gaze heatmap of one participant. Highlighted as white circles and boxes are the regions of interest used for oculomotor analyses. Yellow highlights heatmap for one example participant after cue onset.

**Figure S3.**
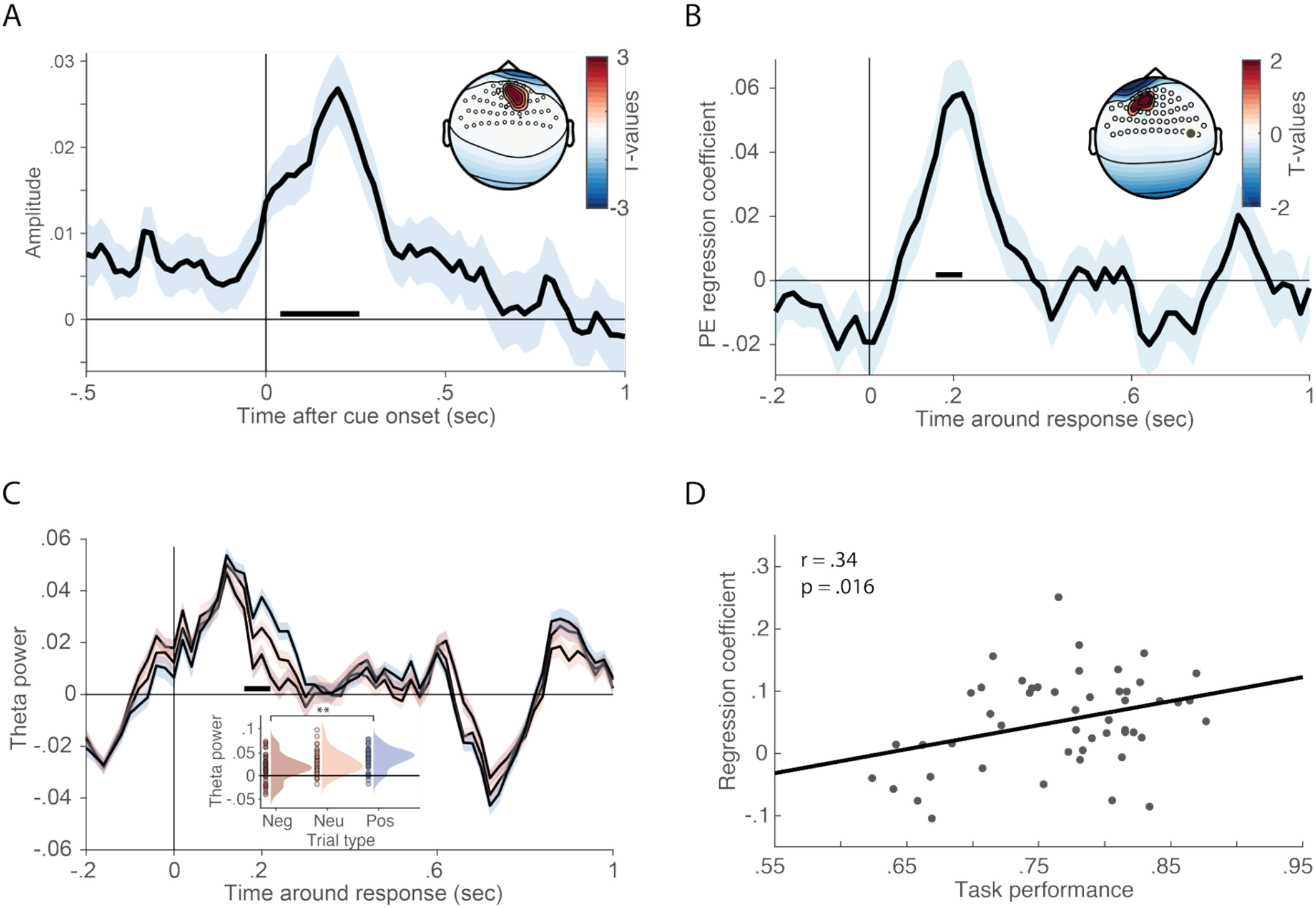
Frontal midline theta associated with prediction error of hidden state inference model. **A**. Greater theta power over midfrontal electrodes on trial 1 compared to subsequent trials, with the two significant clusters averaged for topography. **B**. Model-derived signed prediction error (PE) regressed against frontal midline theta (FMT) revealed a significant cluster approximately 200ms after response onset. **C**. Frontal theta activity as a function of signed prediction error (PE). Trials were divided into participant-specific tertiles based on HSI model-derived signed prediction errors and aligned to the behavioural response. Time courses show mean theta power averaged across the frontocentral cluster identified in the regression analysis, separately for negative, neutral, and positive PE trials. Shaded regions indicate ±SE across participants. The black bar denotes the significant time window identified by the cluster-based permutation test. Inset: Mean theta power averaged across the significant time window. Positive PE trials exhibited significantly greater theta power than negative PE trials (t(49) = 3.39, p = .0014). Points represent individual participants and shaded distributions represent kernel density estimates. **D**. Individual differences in prediction-error signalling predict behavioural performance. The strength of prediction-error-related modulation of frontal theta activity, quantified as the mean regression coefficient within the significant cluster shown in panel B, was positively associated with task accuracy. Participants whose frontal theta responses more strongly tracked model-derived prediction errors tended to perform better on the task. Each point represents a participant; the solid line shows the linear regression fit.

**Figure S4.**
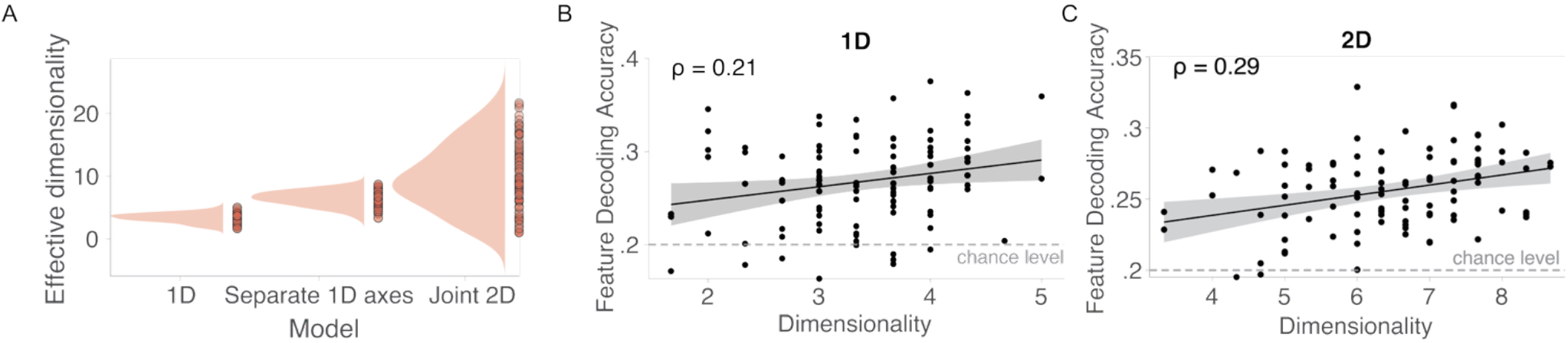
Scaling of neural dimensionality with task space. **A**. Effective dimensionality (ED) for 1D trials, separate 1D axes in 2D trials (calculated as additive ED for the two separate 1D axes) and for joint 2D model in 2D trials. **B**. Partial correlation between decoding accuracy for each level of the attended dimension and ED showed a significant positive correlation for 1D trials. **C**. Same as in (B), but now for 2D trials with separate 1D axes combined, again showing a significant positive correlation, suggesting that an expansion of task space is related to more efficient decoding.

**Figure S5.**
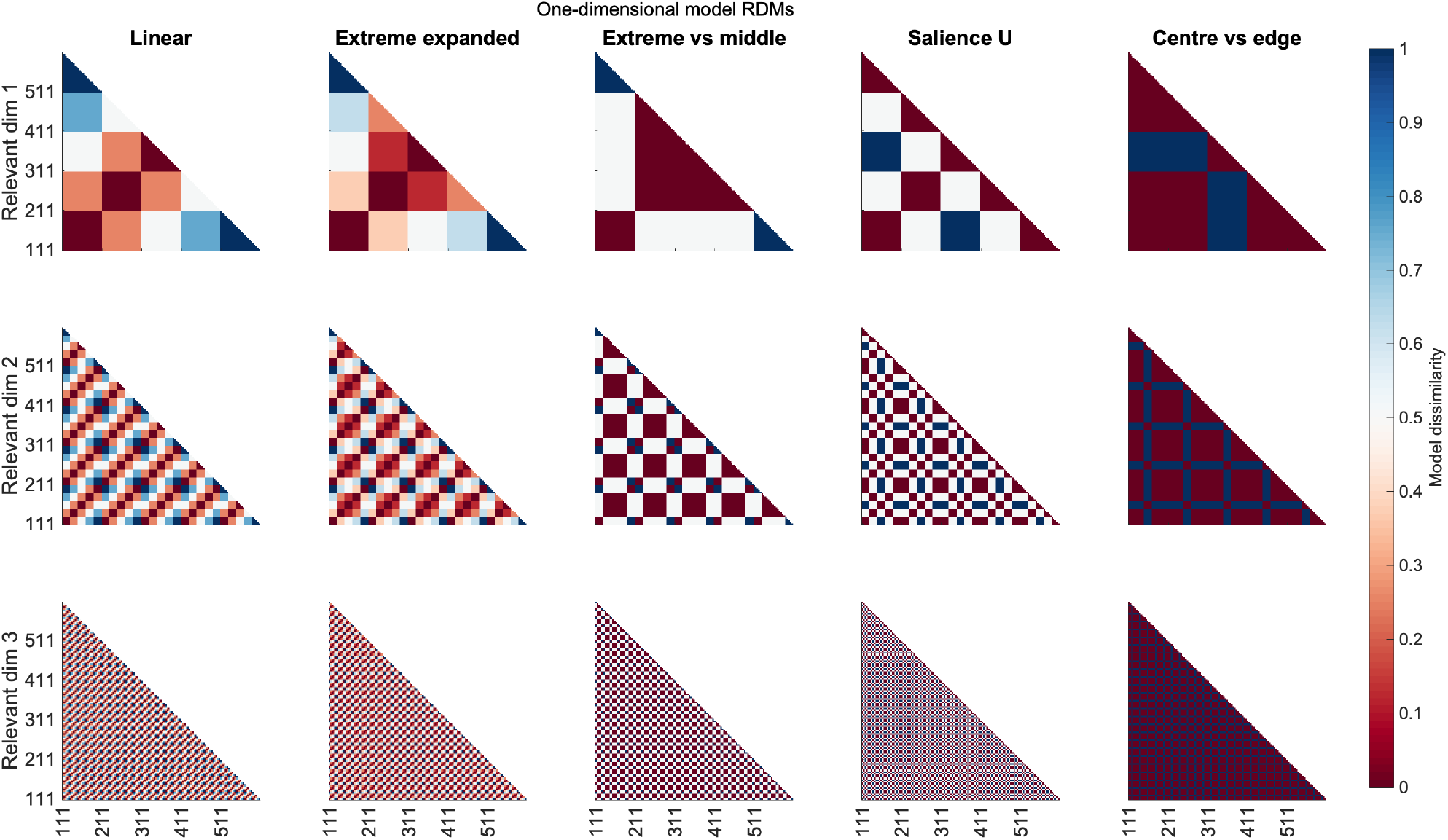
All 1D model representational dissimilarity matrices.

**Figure S6.**
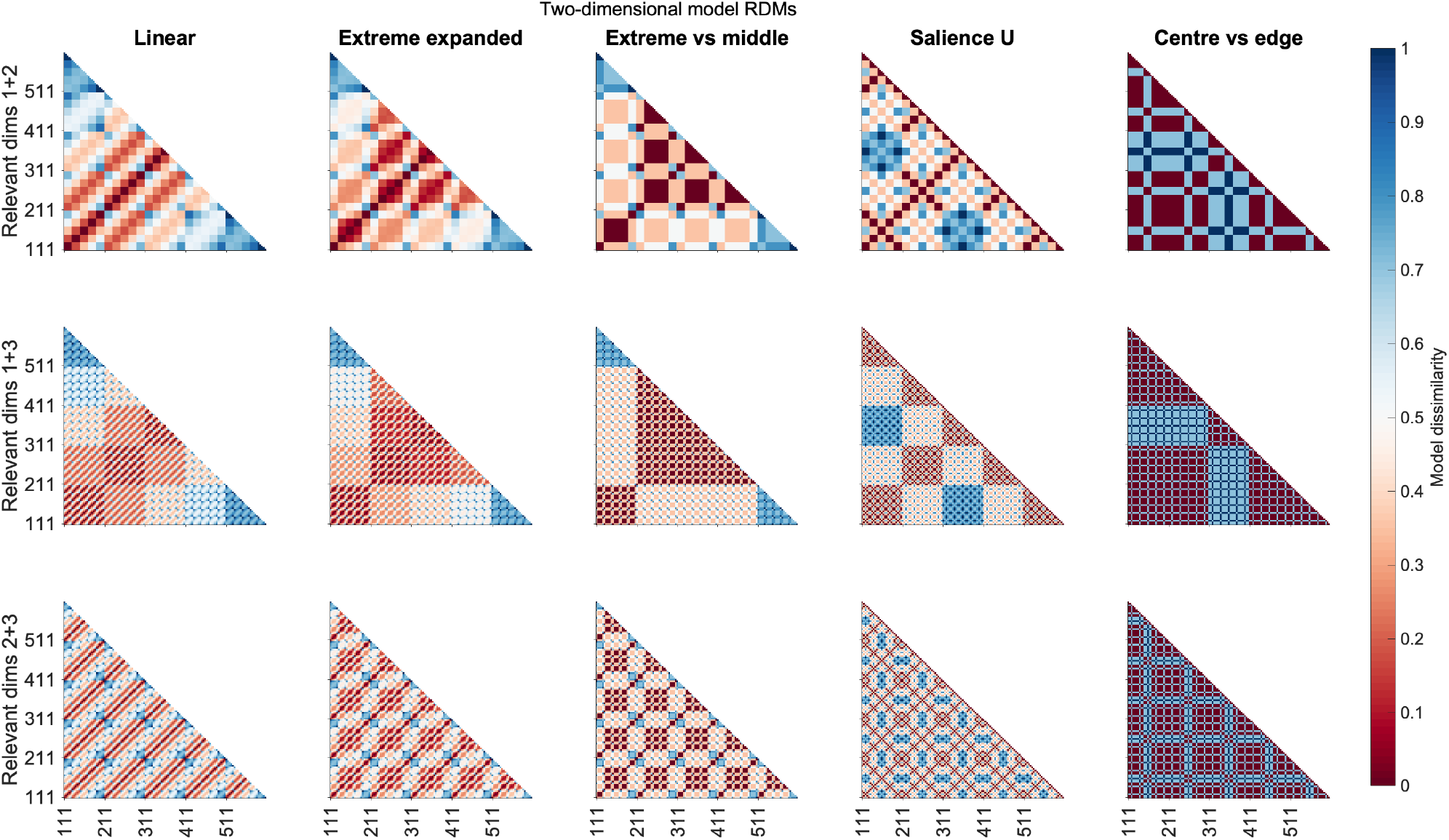
All 2D model representational dissimilarity matrices.

**Table S1.** Parameter values (Mean ± SD) from model fitting for each model.

| Model | Alpha | Alpha nr | Beta | Delta | Beta-action | C | Gamma |
| --- | --- | --- | --- | --- | --- | --- | --- |
| Q | .620 ± .140 | N/A | 2.642 ± .707 | N/A | 14.5899 ± 11.048 | N/A | N/A |
| QF | .273 ± .081 | N/A | 8.965 ± 1.383 | 0.60101 ± 0.27275 | 17.573 ± 9.856 | N/A | N/A |
| QC | .876 ± .177 | .086 ± .118 | 2.379 ± .577 | N/A | 12.344 ± 9.532 | N/A | N/A |
| HSI | N/A | N/A | .487 ± .104 | N/A | 7.3397 ± 2.067 | .838 ± .194 | .288 ± .253 |

## References

Ahlheim, C., & Love, B. C. (2018). Estimating the functional dimensionality of neural representations. NeuroImage, 179, 51–62. 10.1016/j.neuroimage.2018.06.015

Bar-Gad, I., Morris, G., & Bergman, H. (2003). Information processing, dimensionality reduction and reinforcement learning in the basal ganglia. Progress in Neurobiology, 71(6), 439–473. 10.1016/j.pneurobio.2003.12.001

Behrens, T. E. J., Muller, T. H., Whittington, J. C. R., Mark, S., Baram, A. B., Stachenfeld, K. L., & Kurth-Nelson, Z. (2018). What Is a Cognitive Map? Organizing Knowledge for Flexible Behavior. Neuron, 100(2), 490–509. 10.1016/j.neuron.2018.10.002

Bein, O., & Niv, Y. (2025). Schemas, reinforcement learning and the medial prefrontal cortex. Nature Reviews Neuroscience, 26(3), 141–157. 10.1038/s41583-024-00893-z

Bellman, R. (1957). Dynamic Programming. Princeton University Press.

Bellmund, J. L. S., Gärdenfors, P., Moser, E. I., & Doeller, C. F. (2018). Navigating cognition: Spatial codes for human thinking. Science, 362(6415), eaat6766. 10.1126/science.aat6766

Blair, M. R., Watson, M. R., Walshe, R. C., & Maj, F. (2009). Extremely selective attention: Eye-tracking studies of the dynamic allocation of attention to stimulus features in categorization. Journal of Experimental Psychology: Learning, Memory, and Cognition, 35(5), 1196–1206. 10.1037/a0016272

Boyen, X., Friedman, N., & Koller, D. (2013). Discovering the Hidden Structure of Complex Dynamic Systems (arXiv:1301.6683). arXiv. 10.48550/arXiv.1301.6683

Brainard, D. H. (1997). The Psychophysics Toolbox. Spatial Vision, 10, 433–436. 10.1163/156856897X00357

Braun, D. A., Mehring, C., & Wolpert, D. M. (2010). Structure learning in action. Behavioural Brain Research, 206(2), 157–165. 10.1016/j.bbr.2009.08.031

Braunlich, K., & Love, B. C. (2022). Bidirectional influences of information sampling and concept learning. Psychological Review, 129(2), 213–234. 10.1037/rev0000287

Cavanagh, J. F., & Frank, M. J. (2014). Frontal theta as a mechanism for cognitive control. Trends in Cognitive Sciences, 18(8), 414–421. 10.1016/j.tics.2014.04.012

Cavanagh, J. F., & Shackman, A. J. (2015). Frontal midline theta reflects anxiety and cognitive control: Meta-analytic evidence. Journal of Physiology, Paris, 109(1-3), 3–15. 10.1016/j.jphysparis.2014.04.003

Cockburn, J., & Holroyd, C. B. (2018). Feedback information and the reward positivity. International Journal of Psychophysiology, 132, 243–251. 10.1016/j.ijpsycho.2017.11.017

Corbetta, M., & Shulman, G. L. (2002). Control of goal-directed and stimulus-driven attention in the brain. Nature Reviews Neuroscience, 3(3), Article 3. 10.1038/nrn755

Cornelissen, F. W., Peters, E. M., & Palmer, J. (2002). The Eyelink Toolbox: Eye tracking with MATLAB and the Psychophysics Toolbox. Behavior Research Methods, Instruments, & Computers, 34(4), 613–617. 10.3758/BF03195489

Costa, V. D., Tran, V. L., Turchi, J., & Averbeck, B. B. (2015). Reversal learning and dopamine: A bayesian perspective. The Journal of Neuroscience: The Of^icial Journal of the Society for Neuroscience, 35(6), 2407–2416. 10.1523/JNEUROSCI.1989-14.2015

Courellis, H. S., Minxha, J., Cardenas, A. R., Kimmel, D. L., Reed, C. M., Valiante, T. A., Salzman, C. D., Mamelak, A. N., Fusi, S., & Rutishauser, U. (2024). Abstract representations emerge in human hippocampal neurons during inference. Nature, 1–9. 10.1038/s41586-024-07799-x

Daw, N. D., Gershman, S. J., Seymour, B., Dayan, P., & Dolan, R. J. (2011). Model-Based Influences on Humans’ Choices and Striatal Prediction Errors. Neuron, 69(6), 1204–1215. 10.1016/j.neuron.2011.02.027

Duan, Y., Zhan, J., Gross, J., Ince, R. A. A., & Schyns, P. G. (2024). Pre-frontal cortex guides dimension-reducing transformations in the occipito-ventral pathway for categorization behaviors. Current Biology, 34(15), 3392-3404.e5. 10.1016/j.cub.2024.06.050

Edgell, S. E., & Morrissey, J. M. (1987). Delayed exposure to additional relevant information in nonmetric multiplecue probability learning. Organizational Behavior and Human Decision Processes, 40(1), 22–38. 10.1016/0749-5978(87)90003-3

Eichenbaum, H., & Cohen, N. J. (2014). Can We Reconcile the Declarative Memory and Spatial Navigation Views on Hippocampal Function? Neuron, 83(4), 764–770. 10.1016/j.neuron.2014.07.032

Flesch, T., Juechems, K., Dumbalska, T., Saxe, A., & Summerfield, C. (2022). Orthogonal representations for robust context-dependent task performance in brains and neural networks. Neuron, 110(7), 1258-1270.e11. 10.1016/j.neuron.2022.01.005

Frank, M. J., & Badre, D. (2012). Mechanisms of hierarchical reinforcement learning in corticostriatal circuits 1: Computational analysis. Cerebral Cortex (New York, N.Y.: 1991), 22(3), 509–526. 10.1093/cercor/bhr114

Gärdenfors, P. (2000). Conceptual Spaces: The Geometry of Thought. The MIT Press. 10.7551/mitpress/2076.001.0001

Garvert, M. M., Dolan, R. J., & Behrens, T. E. (2017). A map of abstract relational knowledge in the human hippocampal-entorhinal cortex. ELife, 6, e17086. 10.7554/eLife.17086

Gershman, S. J., & Niv, Y. (2010). Learning latent structure: Carving nature at its joints. Current Opinion in Neurobiology, 20(2), 251–256. 10.1016/j.conb.2010.02.008

Gershman, S. J., Norman, K. A., & Niv, Y. (2015). Discovering latent causes in reinforcement learning. Current Opinion in Behavioral Sciences, 5, 43–50. 10.1016/j.cobeha.2015.07.007

Heydari, S., & Holroyd, C. B. (2016). Reward positivity: Reward prediction error or salience prediction error? Psychophysiology, 53(8), 1185–1192. 10.1111/psyp.12673

Jas, M., Engemann, D. A., Bekhti, Y., Raimondo, F., & Gramfort, A. (2017). Autoreject: Automated artifact rejection for MEG and EEG data. NeuroImage, 159, 417–429. 10.1016/j.neuroimage.2017.06.030

Kato, A., & Morita, K. (2016). Forgetting in Reinforcement Learning Links Sustained Dopamine Signals to Motivation. PLoS Computational Biology, 12(10), e1005145. 10.1371/journal.pcbi.1005145

Knudsen, E. B., & Wallis, J. D. (2021). Hippocampal neurons construct a map of an abstract value space. Cell, 184(18), 4640-4650.e10. 10.1016/j.cell.2021.07.010

Konidaris, G. (2019). On the necessity of abstraction. Current Opinion in Behavioral Sciences, 29, 1–7. 10.1016/j.cobeha.2018.11.005

Kowler, E., Anderson, E., Dosher, B., & Blaser, E. (1995). The role of attention in the programming of saccades. Vision Research, 35(13), 1897–1916. 10.1016/0042-6989(94)00279-u

Kruschke, J. K. (1996). Base rates in category learning. Journal of Experimental Psychology: Learning, Memory, and Cognition, 22(1), 3–26. 10.1037/0278-7393.22.1.3

Leong, Y. C., Radulescu, A., Daniel, R., DeWoskin, V., & Niv, Y. (2017). Dynamic Interaction between Reinforcement Learning and Attention in Multidimensional Environments. Neuron, 93(2), 451–463. 10.1016/j.neuron.2016.12.040

Liang, Z., Glitz, L., Hefner, M. B., Lan, D., Klein-Flugge, M., & Summerfield, C. (2025). Distinct roles of hippocampus and neocortex in symbolic compositional generalization. BioRxiv, 2025.08.19.671090. 10.1101/2025.08.19.671090

Lopez-Gamundi, P., Mas-Herrero, E., & Marco-Pallares, J. (2024). Disentangling effort from probability of success: Temporal dynamics of frontal midline theta in effort-based reward processing. Cortex, 176, 94–112. 10.1016/j.cortex.2024.03.014

Love, B. C., Medin, D. L., & Gureckis, T. M. (2004). SUSTAIN: A Network Model of Category Learning. Psychological Review, 111(2), 309–332. 10.1037/0033-295X.111.2.309

Mack, M. L., Preston, A. R., & Love, B. C. (2020). Ventromedial prefrontal cortex compression during concept learning. Nature Communications, 11(1), Article 1. 10.1038/s41467-019-13930-8

Menghi, N., Kacar, K., & Penny, W. (2021). Multitask learning over shared subspaces. PLOS Computational Biology, 17(7), e1009092. 10.1371/journal.pcbi.1009092

Menghi, N., Silvestrin, F., Pascolini, L., & Penny, W. (2023). The emergence of task-relevant representations in a nonlinear decision-making task. Neurobiology of Learning and Memory, 206, 107860. 10.1016/j.nlm.2023.107860

Mishchanchuk, K., Gregoriou, G., Qü, A., Kastler, A., Huys, Q. J. M., Wilbrecht, L., & MacAskill, A. F. (2024). Hidden state inference requires abstract contextual representations in the ventral hippocampus. Science, 386(6724), 926–932. 10.1126/science.adq5874

Moore, T., & Fallah, M. (2001). Control of eye movements and spatial attention. Proceedings of the National Academy of Sciences of the United States of America, 98(3), 1273–1276. 10.1073/pnas.98.3.1273

Najemnik, J., & Geisler, W. S. (2005). Optimal eye movement strategies in visual search. Nature, 434(7031), 387–391. 10.1038/nature03390

Nelson, J. D., & Cottrell, G. W. (2007). A probabilistic model of eye movements in concept formation. Neurocomputing, 70(13), 2256–2272. 10.1016/j.neucom.2006.02.026

Niv, Y. (2019). Learning task-state representations. Nature Neuroscience, 22(10), 1544–1553. 10.1038/s41593-019-0470-8

Niv, Y., Daniel, R., Geana, A., Gershman, S. J., Leong, Y. C., Radulescu, A., & Wilson, R. C. (2015). Reinforcement Learning in Multidimensional Environments Relies on Attention Mechanisms. The Journal of Neuroscience, 35(21), 8145–8157. 10.1523/JNEUROSCI.2978-14.2015

Nosofsky, R. M. (1986). Attention, similarity, and the identification-categorization relationship. Journal of Experimental Psychology: General, 115, 39–57. 10.1037/0096-3445.115.1.39

Oostenveld, R., Fries, P., Maris, E., & Schoffelen, J.-M. (2011). FieldTrip: Open source software for advanced analysis of MEG, EEG, and invasive electrophysiological data. Computational Intelligence and Neuroscience, 2011, 156869. 10.1155/2011/156869

Palminteri, S., Khamassi, M., Joffily, M., & Coricelli, G. (2015). Contextual modulation of value signals in reward and punishment learning. Nature Communications, 6(1), 8096. 10.1038/ncomms9096

Park, S. A., Miller, D. S., Nili, H., Ranganath, C., & Boorman, E. D. (2020). Map Making: Constructing, Combining, and Inferring on Abstract Cognitive Maps. Neuron, 107(6), 1226-1238.e8. 10.1016/j.neuron.2020.06.030

Park, S. A., Zolfaghar, M., Russin, J., Miller, D. S., O’Reilly, R. C., & Boorman, E. D. (2023). The representational geometry of cognitive maps under dynamic cognitive control (p. 2023.02.04.527142). bioRxiv. 10.1101/2023.02.04.527142

Radulescu, A., Niv, Y., & Ballard, I. (2019). Holistic Reinforcement Learning: The Role of Structure and Attention. Trends in Cognitive Sciences, 23(4), 278–292. 10.1016/j.tics.2019.01.010

Rehder, B., & Hoffman, A. B. (2005). Eyetracking and selective attention in category learning. Cognitive Psychology, 51(1), 1–41. 10.1016/j.cogpsych.2004.11.001

Schuck, N. W., Cai, M. B., Wilson, R. C., & Niv, Y. (2016). Human Orbitofrontal Cortex Represents a Cognitive Map of State Space. Neuron, 91(6), 1402–1412. 10.1016/j.neuron.2016.08.019

Shiferaw, B., Downey, L., & Crewther, D. (2019). A review of gaze entropy as a measure of visual scanning efficiency. Neuroscience and Biobehavioral Reviews, 96, 353–366. 10.1016/j.neubiorev.2018.12.007

Starkweather, C. K., Gershman, S. J., & Uchida, N. (2018). The Medial Prefrontal Cortex Shapes Dopamine Reward Prediction Errors under State Uncertainty. Neuron, 98(3), 616-629.e6. 10.1016/j.neuron.2018.03.036

Sutton, R. S., & Barto, A. G. (2018). Reinforcement learning: An introduction, 2nd ed (pp. xxii, 526). The MIT Press.

Theves, S., Fernández, G., & Doeller, C. F. (2020). The Hippocampus Maps Concept Space, Not Feature Space. The Journal of Neuroscience, 40(38), 7318–7325. 10.1523/JNEUROSCI.0494-20.2020

Vertechi, P., Lottem, E., Sarra, D., Godinho, B., Treves, I., Quendera, T., Oude Lohuis, M. N., & Mainen, Z. F. (2020). Inference-Based Decisions in a Hidden State Foraging Task: Differential Contributions of Prefrontal Cortical Areas. Neuron, 106(1), 166-176.e6. 10.1016/j.neuron.2020.01.017

Watkins, C. J. C. H., & Dayan, P. (1992). Q-learning. Machine Learning, 8(3), 279–292. 10.1007/BF00992698

Wilson, R. C., & Collins, A. G. (2019). Ten simple rules for the computational modeling of behavioral data. ELife, 8, e49547. 10.7554/eLife.49547

Wilson, R. C., & Niv, Y. (2012). Inferring Relevance in a Changing World. Frontiers in Human Neuroscience, 5, 189. 10.3389/fnhum.2011.00189

Wilson, R. C., Takahashi, Y. K., Schoenbaum, G., & Niv, Y. (2014). Orbitofrontal cortex as a cognitive map of task space. Neuron, 81(2), 267–279. 10.1016/j.neuron.2013.11.005

Yang, S. C.-H., Lengyel, M., & Wolpert, D. M. (2016). Active sensing in the categorization of visual patterns. ELife, 5, e12215. 10.7554/eLife.12215

Zhang, M., & Yu, Q. (2024). The representation of abstract goals in working memory is supported by task-congruent neural geometry. PLoS Biology, 22(12), e3002461. 10.1371/journal.pbio.3002461

Zhang, X.-Y., Bobadilla-Suarez, S., Luo, X., Lemonari, M., Brincat, S. L., Siegel, M., Miller, E. K., & Love, B. C. (2023). Adaptive stretching of representations across brain regions and deep learning model layers [Preprint]. Neuroscience. 10.1101/2023.12.01.569615

